# Ethylene and ROS Signaling Are Key Regulators of Lateral Root Development under Salt Stress in Tomato

**DOI:** 10.1101/2024.06.20.599848

**Authors:** Maryam Rahmati Ishka, Jiantao Zhao, Hayley Sussman, Devasantosh Mohanty, Li’ang Yu, Eric Craft, Miguel Pineros, Mark Tester, Dorota Kawa, Ron Mittler, Andrew Nelson, Zhangjun Fei, Magdalena Julkowska

**Affiliations:** Boyce Thompson Institute, Ithaca, NY, USA; Division of Plant Science and Technology, College of Agriculture Food and Natural Resources, Christopher S. Bond Life Sciences Center, 1201 Rollins St., University of Missouri, Columbia, MO 65211, USA; USDA-ARS, Ithaca, NY, USA; Center for Desert Agriculture, King Abdullah University of Science and Technology, Thuwal, Saudi Arabia; Environmental and Computational Plant Development & Plant Stress Resilience, Utrecht University, Utrecht, The Netherlands

**Author notes:** Maryam Rahmati Ishka -, Jiantao Zhao -, Hayley Sussman -, Devasantosh Mohanty -, Li’ang Yu -, Eric Craft -, Miguel Pineros -, Mark Tester -, Dorota Kawa -, Ron Mittler -, Andrew Nelson -, Zhangjun Fei.

## Abstract

Roots are the primary site for sensing salt, making them critical to understanding how plants adapt to salinity. Unraveling the genetic and molecular mechanisms that maintain root growth under salt stress is essential for developing resilient crops. Tomatoes are vulnerable to salinity, as over 50% of arable land is projected to become saline by 2050. While salt stress effects on tomato seed germination, shoot growth, and fruit yield are well-documented, the genetic basis of root development under salinity remains underexplored. Previous studies focused on physiological responses of roots in a narrow range of cultivated-tomatoes, overlooking the genetic diversity for salt resilience in wild-relatives like *Solanum pimpinellifolium*. Here, we investigated salt-induced changes in root system architecture (RSA) across a natural diversity panel of 220 wild- and 25 cultivated-tomato varieties. We identified tolerant accessions with different RSA strategies; prioritizing lateral root elongation versus emergence. To identify genes involved in specific aspects of lateral root development, an F1 hybrid was generated that exhibited both parents’ combined characteristics. An F2-segregating population was then used to identify four distinct subpopulations for Bulk Segregant Analysis (BSA). Simultaneously, we conducted Genome-Wide-Association Studies (GWAS) on root architecture of wild-tomato accessions under salt stress. By integrating the results of BSA and GWAS, we identified 22 candidates involved in the maintenance of root architecture. We utilized RNA-Seq to examine transcriptome reprogramming of the 22 genes in tomato accessions with contrasting lateral root responses. This approach resulted in the identification of 2 genes, AP2-like-ethylene-responsive transcription factor TOE3 and L-ascorbate peroxidase. Using exogenous ethylene, we demonstrated that root architecture exhibits plasticity, which benefits the plant by reducing Na^+^ accumulation. We profiled H_2_O_2_ waves across root systems of tomatoes, emphasizing the role of ROS in maintaining root system architecture under salt stress. These findings provide novel genetic targets for enhancing salt resilience in tomatoes, opening avenues for future research and breeding programs.

## Introduction

Tomatoes are one of the most widely cultivated and economically significant crops globally, contributing substantially to food security and agricultural economies. With projections indicating that over 50% of arable land will become saline by 2050, understanding tomato responses to diverse saline environments becomes paramount (Roșca et al. 2023). In arid regions, where evaporation rates exceed freshwater input, soil salinization is contentiously increasing (Richards 1969; Abrol et al. 1988; Fischer et al. 2021). These dryland areas, including regions in South America, Southern and Western Australia, Mexico, Southwest United States, and South Africa, have been identified as hotspots for soil salinization (Hassani et al. 2021). Additionally, salt interactions with clay particles result in impenetrable claypan that prevents roots from reaching deeper soil levels (Cione et al. 2000).

Roots are the primary site of salinity sensing and serve to integrate salt stress with other environmental factors, such as nutrient and water availability. The susceptibility of roots to salt stress determines the plant’s overall productivity (Atkin et al. 1973; Steppuhn and Raney 2005). Therefore, understanding the molecular processes leading to maintained root growth is necessary for enhancing crop environmental resilience. Although the effects of salt stress on tomato plants have been studied in the context of seed germination, shoot growth, and fruit yield (Manaa et al. 2011; Yang et al. 2019; Sousa et al. 2022; Morton et al. 2023), the impact of salt stress on root architecture remains unexplored. Furthermore, previous studies have primarily focused on a limited number of cultivars and assessed physiological responses (Manaa et al. 2011), rather than elucidating the genetic basis of salt resilience. As freshwater scarcity intensifies, unraveling root-driven salt stress resilience is key to boosting global agricultural productivity and sustainability (Asins et al. 2010; Melino and Tester 2023).

Wild relatives of crops possess valuable traits such as disease resistance, stress tolerance, and nutritional quality, which can be introduced into cultivated varieties through breeding programs (Dempewolf et al. 2014; Ebert and Schafleitner 2015; Brozynska et al. 2016; Pailles et al. 2020; Li et al. 2023). Understanding the genetic basis of these traits in wild relatives facilitates the development of more resilient crop varieties, which are crucial for ensuring food security in the face of environmental challenges. The closest wild relative of cultivated tomato, *Solanum pimpinellifollium* (Pease et al. 2016), shows high tolerance to many biotic and abiotic stresses (Yassin 1985; Sarfatti et al. 1991; Paran and van der Knaap 2007; Ashrafi et al. 2009; Celik et al. 2017; Zhang et al. 2018). However, only a few studies explored the genetics underlying salinity tolerance within *S. pimpinellifolium* natural diversity panels (Rao et al. 2015; Wang et al. 2020b; Morton et al. 2023).

Here, we investigated salt-induced changes in root architecture across a natural diversity panel of 220 wild- and 25 cultivated-tomato varieties. We conducted Genome-Wide-Association Studies (GWAS) on root architecture of wild-tomato accessions under salt stress and identified 55 putative candidate loci. We identified tolerant accessions with different lateral root characteristics, prioritizing either lateral root elongation or emergence. By crossing the tolerant accessions, we generated F1 hybrids that exhibited both parents’ combined characteristics. We developed an F2-segregating population and identified four distinct subpopulations for Bulk Segregant Analysis (BSA). By integrating the results of BSA and GWAS, we identified 22 candidate genes involved in the maintenance of root architecture. We utilized RNA-Seq to examine transcriptome reprogramming of the 22 genes in tomato accessions with contrasting lateral root responses. This approach enabled identification of four candidate genes, two with existing annotation including AP2-like-ethylene-responsive transcription factor TOE3 and L-ascorbate peroxidase. Using exogenous ethylene, we demonstrated that root architecture exhibits plasticity, which benefits the plant by reducing Na^+^ accumulation. We profiled H_2_O_2_ accumulation across root systems of tomatoes under salt stress, revealed a unique ROS accumulation pattern associated with maintaining root system architecture of different accessions under salt stress. These findings provide novel genetic targets for enhancing salt resilience in tomatoes, opening avenues for future research and breeding programs.

## Results

### Wild tomato shows high variation in lateral root development in response to salt stress

Using an agar plate system, we evaluated 220 *S. pimpinellifolium* accessions for their responses to salt stress (**Table S1**). The 4-day-old seedlings were transferred to media containing 0 or 100 mM NaCl and scanned daily. Although the images were collected for one week after transfer, the root architecture was quantified exclusively for the first 4 days after transfer due to root architecture complexity in a plate setup. We have calculated the daily Main Root growth rate (**Fig. 1A, Fig. S1**), increase in Lateral Root number (**Fig. 1B**), and increase in average Lateral Root Length (**Fig. 1C**) using linear functions. Although salt treatment only slightly decreased the Main Root Growth Rate from 1.334 to 1.275 cm / day (**Fig. 1A**), there was a significant effect of both treatment (ANOVA p-value of 0.000157) and genotype (ANOVA p-value < 2-e16), and interaction between genotype (G) x treatment (T) (**Fig. 1D**, ANOVA *GxT* p-value of 9.56e-16). On the other hand, Lateral Root emergence was substantially decreased by salt stress (from 4.482 to 2.219 LR / day). This interaction was significantly affected by genotype (**Fig. 1 E**, ANOVA *GxT* p-value of 3.59e-14). Similar to the Main Root growth rate, the Lateral Root growth rate showed little decrease in response to salt stress (from 0.315 to 0.304 cm / day), and this phenotype was exclusively affected by genotype (ANOVA p-value < 2e-16), but not by environment (ANOVA p-value of 0.188) or interaction between genotype and environment (**Fig. 1F**, ANOVA *GxT* p-value of 0.383). To evaluate the extent to which root architecture is reprogrammed in response to salt stress, meaning whether individual root architecture components are affected differently, we calculated stress tolerance indices (STI = GR at 100 mM / GR at 0 mM NaCl, **Fig. 1G**). We observed significant differences between individual root architecture components, with the increase in lateral root number being affected the most (0.494 of the control values) and average lateral root growth being affected the least (1.034 of the control value). Additionally, we observed a significant interaction between the root architecture component and the genotype (Kruskal-Wallis p-value < 2.2e-16), indicating that the tomato root architecture is substantially reprogrammed in response to salt stress. Within the root architecture patterns in the entire diversity panel, we observed that the genotypes that tend to maintain a high number of lateral root numbers, tend to be compromised in their lateral root elongation, while the genotypes that tend to maintain a high lateral root elongation are oftentimes compromised in their lateral root emergence (**Fig. 1G**). These results suggest that the natural variation in *S. pimpinellifolium*’s root architecture in response to salt stress is primarily caused by its differential sensitivity to salt stress in lateral root development.

**Figure 1.**
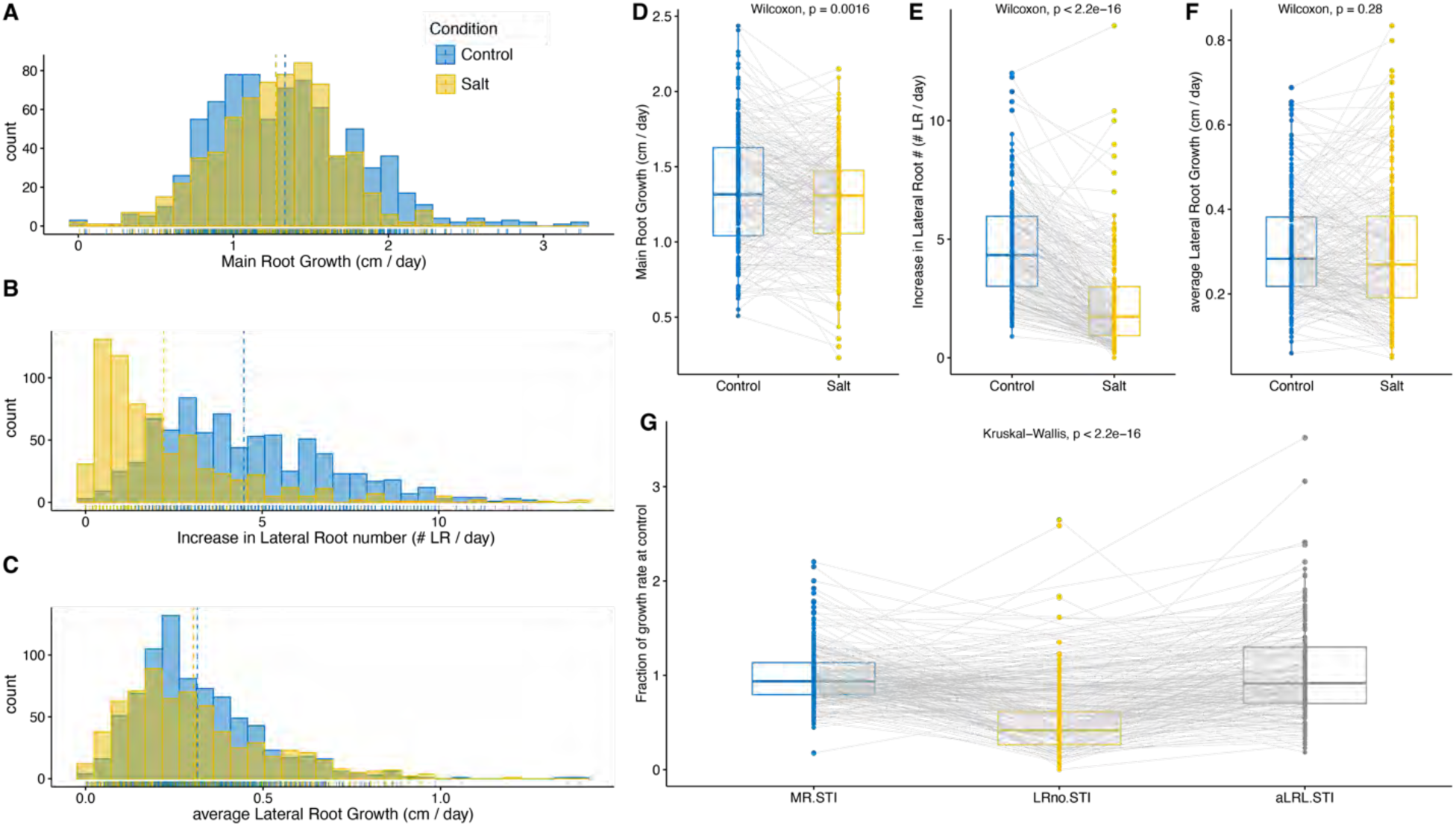
Diversity of salt stress responses observed within the *S. pimpinellifolium* diversity panel. The 220 genotypes of *S. pimpinellifolium* were germinated on ½ MS agar plates for 4 days and subsequently transferred to media containing 0 or 100 mM NaCl (Control and Salt respectively). The seedlings were imaged every 24 h after transfer for 7 consecutive days. The Root Architecture of seedlings was quantified for the first 4 days after transfer, and the **(A)** growth of Main Root, **(B)** increase in Lateral Root number and **(C)** growth of average Lateral Root was estimated using linear growth functions. The population average values for Control and Salt conditions are indicated using the dashed lines. The effect of the treatment was evaluated using a paired Wilcoxon test for **(D)** Main Root Growth Rate, **(E)** increase in Lateral Root number and **(F)** average Lateral Root Growth Rate. The individual gray lines illustrate the genotype-specific change in individual growth parameters across the conditions. **(G)** The genotype-specific growth rates at individual conditions were used to calculate the Stress Tolerance Index (STI) for individual traits, calculated by dividing growth rate estimated under Control / Salt conditions. The relative change in STI represents the salt-induced reprogramming of the root architecture. The individual gray lines represent the genotype-specific change in STI across individual components of root architecture. The individual dots represent the genotype-specific mean calculated from 4 biological replicates.

### Identification of associations with tomato root architecture through GWAS

To identify genetic components underlying the root system architecture responses to salt stress, the collected phenotypic data were combined with previously developed genotypic data (Morton et al. 2023) and used in Genome Wide Association Study (GWAS). Initially, we removed heterozygous markers, imputed the missing markers, and performed GWAS using ASReml script, similarly to the method described in (Awlia et al. 2021). This method yielded a total of 109 associated SNPs above Bonferroni threshold (LOD > 7.15), with 85 unique SNPs, grouping into 16 individual loci out of which 5 loci were associated exclusively with root architecture measured under salt stress (**Table S2**). However, this method required us to use a heavily reduced set of SNPs (708 545). Therefore, we explored additional associations using an entire set of 17 962 579 SNP using the EMMAX model with corrections for population structure (using three PCA) and kinship matrix (Kang et al. 2010). Although no overlap was found between ASReml and EMMA-X based scripts, we identified additional 82 associations above the Bonferroni threshold (> 8.56) with various root system architecture components (**Table S2**). Only one of the significantly associated SNPs had a minor allele frequency (MAF) > 1% (chromosome 1, position 1369582), associated with lateral root distribution over the length of main root (calculated as Center of Gravity) for roots grown under control conditions (**Table S2**). We further inspected associations with LOD score above 5, considering MAF, and the number of SNPs within the 10 kbp interval to select associations that can be pursued further. We identified 55 genetic loci, each containing at least 3 SNPs within 10 kbp interval or mapped/associated with multiple root architecture traits (**Table S2**). Additionally, we adjusted the Bonferroni threshold to 6.77 as the number of SNPs with MAF > 5% was reduced to 5 944 147.

The highest-ranking loci for biomass distribution between main and lateral roots in the first two days (MRLpTRS) were identified on chromosomes 3 and 4, with one locus on chromosome 3 (29 SNPs) and three on chromosome 4 (240, 39, and 85 SNPs) (**Fig. S2A-B**). The chromosome 3 locus mapped to Spimp03g102810, encoding an ABC transporter A family member 7 (**Fig. S2C**). On chromosome 4, the loci spanned a region containing multiple functionally diverse genes, including DUF4283 proteins, L-ascorbate oxidases, MADS-box proteins, RING domain-containing proteins, and a Zinc knuckle CCHC-type family protein (**Fig. S2D-F**). For lateral root length and number after one day of salt stress, high-ranking loci were found on chromosomes 1, 3, and 11 (**Fig. S3A-B**). On chromosome 1, 27 SNPs were located in the promoter region of Spimp01g004220, encoding a LITAF domain-containing protein (**Fig. S3C**). Three SNPs on chromosome 3 spanned genes involved in glycosylation, protein transport, and tRNA synthesis (**Fig. S3D**). The chromosome 11 locus (60 SNPs) was mapped to Spimp11g311320, encoding a chloroplast ribosome-recycling factor, alongside neighboring genes with roles in storage and redox regulation (**Fig. S3E**). Additional loci were linked to increased lateral root number and main root elongation under salt stress (**Fig. S4A-B**). On chromosome 12, 16 SNPs were associated with Spimp12g340040, encoding a glucan endo-1,3-beta-D-glucosidase (**Fig. S4C**). Another 15 SNPs on chromosome 11 were mapped to genes encoding two unknown proteins and a dephospho-CoA kinase (**Fig. S4D**). Other loci with more than 10 highly associated SNPs were identified on chromosomes 1, 2, 5, and 12, linked to root traits under control and salt stress conditions (**Fig. S5A-I**).

### LA2540 and LA1511 prioritize different aspects of lateral root development under salt stress

Based on the root system architecture responses to salt stress observed in the diversity panel, we selected two accessions for further detailed investigation. We selected LA2540 as wild tolerant (referred hereafter as wild tolerant) accession, and LA1371 as wild sensitive (referred hereafter as wild sensitive) accession. Additionally, we included an *S. lycopersicum* accession (LA1511, referred hereafter as cultivated tomato), based on the root system architecture responses to salt stress observed in the cultivated diversity panel and its overall similarity in plant morphology to *S. pimpinellifolium* observed in the greenhouse conditions (e.g. plant and fruit size). Salt stress reduced main root growth (**Fig. 2A**), lateral root length (**Fig. 2B**), and the number of lateral roots (**Fig. 2C**) across all examined accessions. While the STI of the main root did not differ across examined genotypes (**Fig. 2D**), substantial differences were observed between wild-tolerant and cultivated tomatoes in lateral root-related traits (**Fig. 2E-F**). Wild-tolerant accession showed a higher STI for average lateral root elongation (**Fig. 2E**), whereas cultivated tomatoes exhibited a higher STI for lateral root emergence (**Fig. 2F**). Additionally, we evaluated the selected genotypes for ion accumulation using ICP-AES after 10 days of exposure to salt stress (**Fig. 2G and Fig. S6**). Salt stress exposure resulted in increased and decreased Na^+^ and K^+^ accumulation respectively, with no significant differences between the genotypes. Although all accessions maintained a similar Na^+^/K^+^ ratio in the roots, cultivated tomatoes exhibited a significantly lower Na^+^/K^+^ ratio in the shoot compared to both wild accessions. In summary, we have identified LA1511 as a salt-tolerant *S. lycopersicum* line that prioritizes the development of new lateral roots, while LA2540 as a tolerant *S. pimpinellifolium* line that prioritizes the elongation of existing lateral roots.

**Figure 2.**
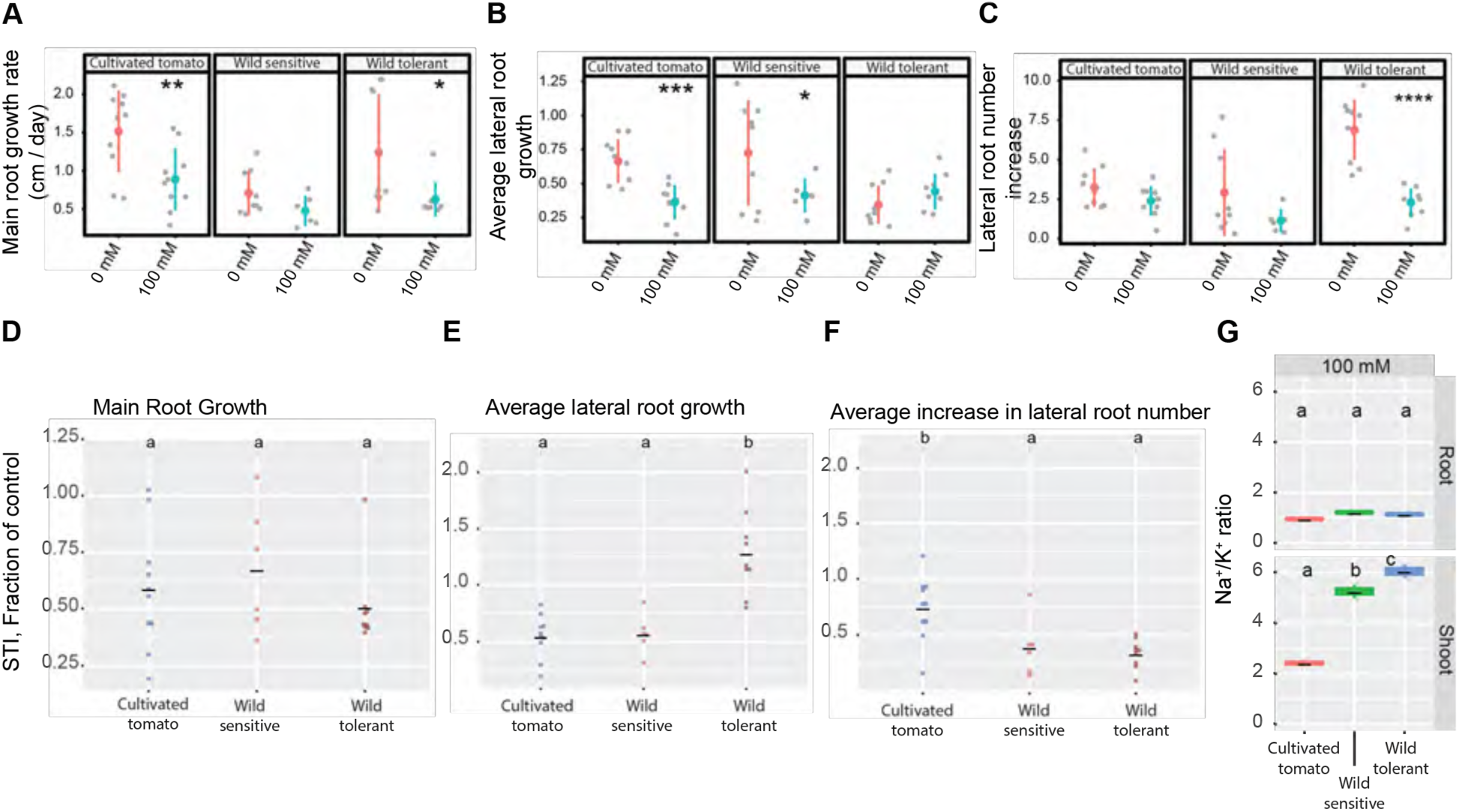
Wild tolerant and cultivated tomatoes have different salt tolerant features regarding lateral root development under the salt. **(A-C)** Root system architecture analysis of cultivated and wild tomatoes under 0 or 100 mM concentrations of NaCl are shown**. (D)** Salt tolerance indexes (STIs) are shown for the main root, **(E)** average lateral root length, **(F)** and lateral root number. The STI was calculated by dividing the growth rate measured under salt stress by the growth rate measured under control condition for each accession. **(G)** Na^+^/K^+^ ratio in root and shoot of different accessions after 10 days on treatment plates. Each dot represents an individual replicate per accession. Lines in (A-C) graphs represent median values. The asterisks above the graphs in (A-C) indicate significant differences between control and salt stress conditions, as determined by the Student ‘s t-test: *P < 0.05, **P < 0.01, ***P < 0.001, and ****P < 0.0001. Statistical analysis was done by comparison of the means for all pairs using Tukey HSD test for (D-G). Levels not connected by the same letter are significantly different (P < 0.05).

### Compound effect of F1 hybrid on maintenance of lateral root development under salt stress

To further evaluate the genetic mechanisms underlying the differential responses of lateral roots between wild tolerant and cultivated tomato and explore whether the salt tolerance mechanisms are complementary between these two accessions, we generated an F1 hybrid by crossing cultivated and wild-tolerant tomato accessions (**Fig. 3**). We investigated root system architecture of F1 individuals along with their parental lines and wild sensitive accession, under non-stress and salt stress conditions (**Fig. 3 and Fig. S7**). While no significant differences were observed in the main root growth rate between the examined accessions under control conditions (**Fig. 3B**), F1 displayed a significant increase in main root growth rate under the salt stress compared to the other three genotypes, especially when compared to both wild tomatoes (**Fig. 3B**). F1 and cultivated tomatoes showed an increase in lateral root growth rate compared to two wild tomatoes under control conditions (**Fig. 3C**). Under salt stress, F1 individuals exhibited a remarkable increase in lateral root growth rate compared to all other accessions (**Fig. 3C**). Regarding increase in lateral root number, F1 and wild-tolerant accession exhibited higher rate of lateral root emergence compared to cultivated and wild-sensitive accessions under control condition (**Fig. 3D**), while no significant differences were observed for absolute emergence rates under salt stress (**Fig. 3D**).

**Figure 3.**
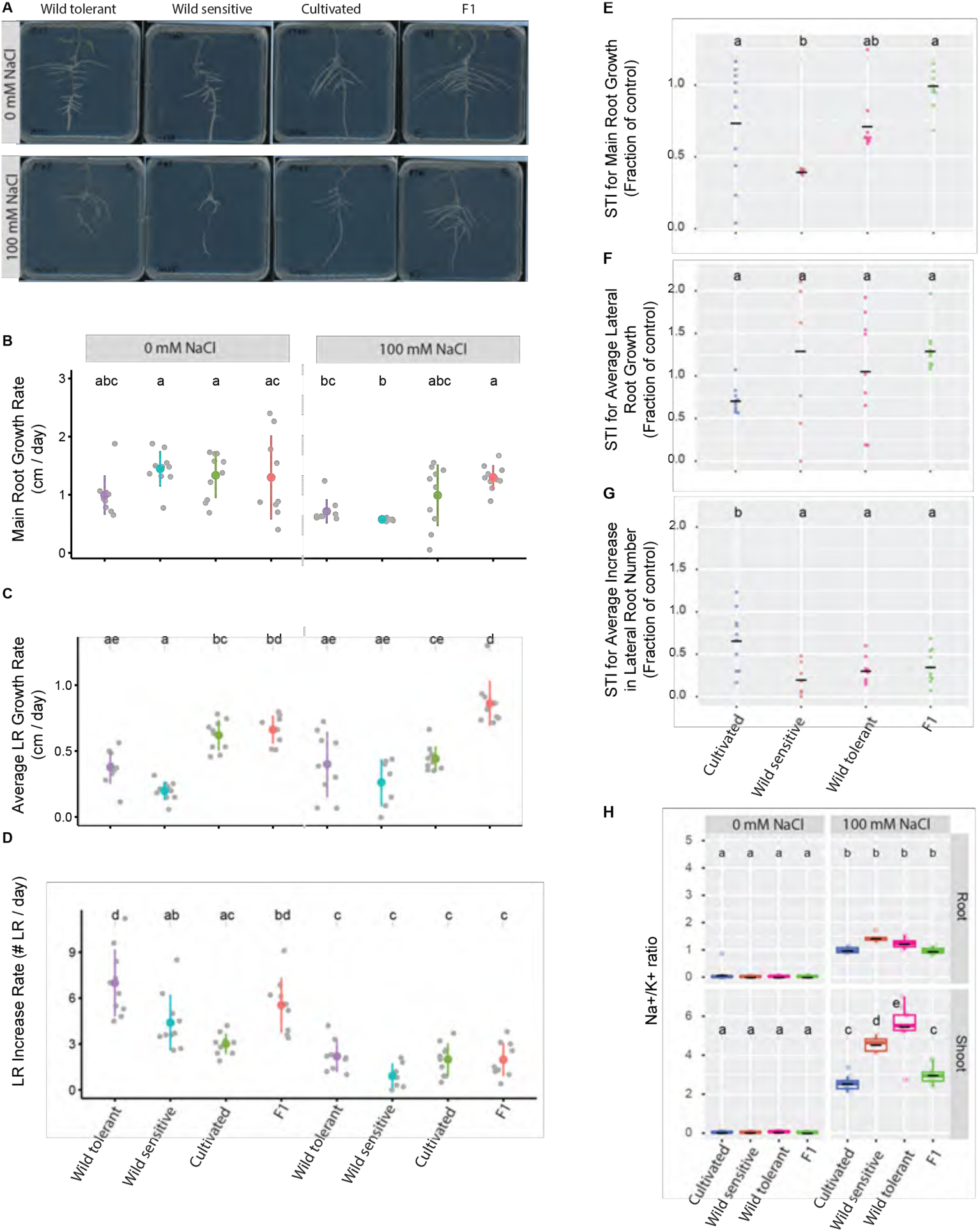
F1 individuals display an increase in main root growth under salt while maintaining lower Na^+^/K^+^ ratio in root and shoot. Root system architecture of F1 individuals, their corresponding parents, i.e., cultivated and wild tolerant tomatoes along with wild sensitive tomatoes were investigated with and without 100 mM NaCl. **(A)** Shown are representative images of 9-day-old seedlings that experienced 5 days of salt treatment. Growth rate for **(B)** main root length**, (C)** lateral root length, **(D)** and lateral root number. Salt tolerance indexes (STIs) are shown for the **(E)** main root, **(F)** average lateral root length, and **(G)** lateral root number. The STI was calculated by dividing the growth rate measured under salt stress by the growth rate measured under control condition for each accession. **(H)** Na^+^/K^+^ ratio of root and shoot of different accessions after 10 days on treatment plates. Each dot represents individual replicas per accession. Statistical analysis was done by comparison of the means for all pairs using Tukey HSD test in all graphs. Levels not connected by the same letter are significantly different (P < 0.05).

To investigate how salt stress impacted various aspects of root architecture, we calculated the STI for main root growth, lateral root elongation, and increased lateral root number (**Fig. 3E-G**). F1 showed the highest STI for main root and lateral root growth rates, compared to its parental lines, indicating a synergistic interaction on these phenotypes. On the other hand, F1 inherited the STI for lateral root emergence, similar to wild tomatoes. We further evaluated F1 along with their parental lines for Na^+^ and K^+^ accumulation and observed that F1 seedlings maintained a low Na^+^/K^+^ ratio in the root and shoot, similar to the cultivated tomato (**Fig. 3H, Fig. S7**). Together, these results indicate that the salt tolerance traits, including maintenance of lateral root development and sodium exclusion, are dominant traits inherited from the tolerant parent. Additionally, F1 displays synergistic effect (hybrid vigor) for main and lateral root growth rates.

### Segregating F2 population reveals loci on chromosomes 3, 6 and 9 to be associated with lateral root development

To identify genomic regions associated with root system architectural traits under salt stress, we generated the F2 population. We investigated the root system architecture of 180 F2 individuals alongside their parental lines under salt stress and identified four phenotypic sub-populations that segregated into four classes regarding their lateral root phenotypes (**Fig. 4A**). The classes comprised individuals with low lateral root length/low lateral root number (low LRL/low LR.no), high lateral root length/low lateral root number (high LRL/low LR.no), low lateral root length/high lateral root number (low LRL/high LR.no), and high lateral root length/ high lateral root number (high LRL/high LR.no). We pooled 25 seedlings from each sub-population, extracted DNA, and performed whole-genome sequencing in the pools and their two parents for Bulk Segregant Analysis (BSA).

**Figure 4.**
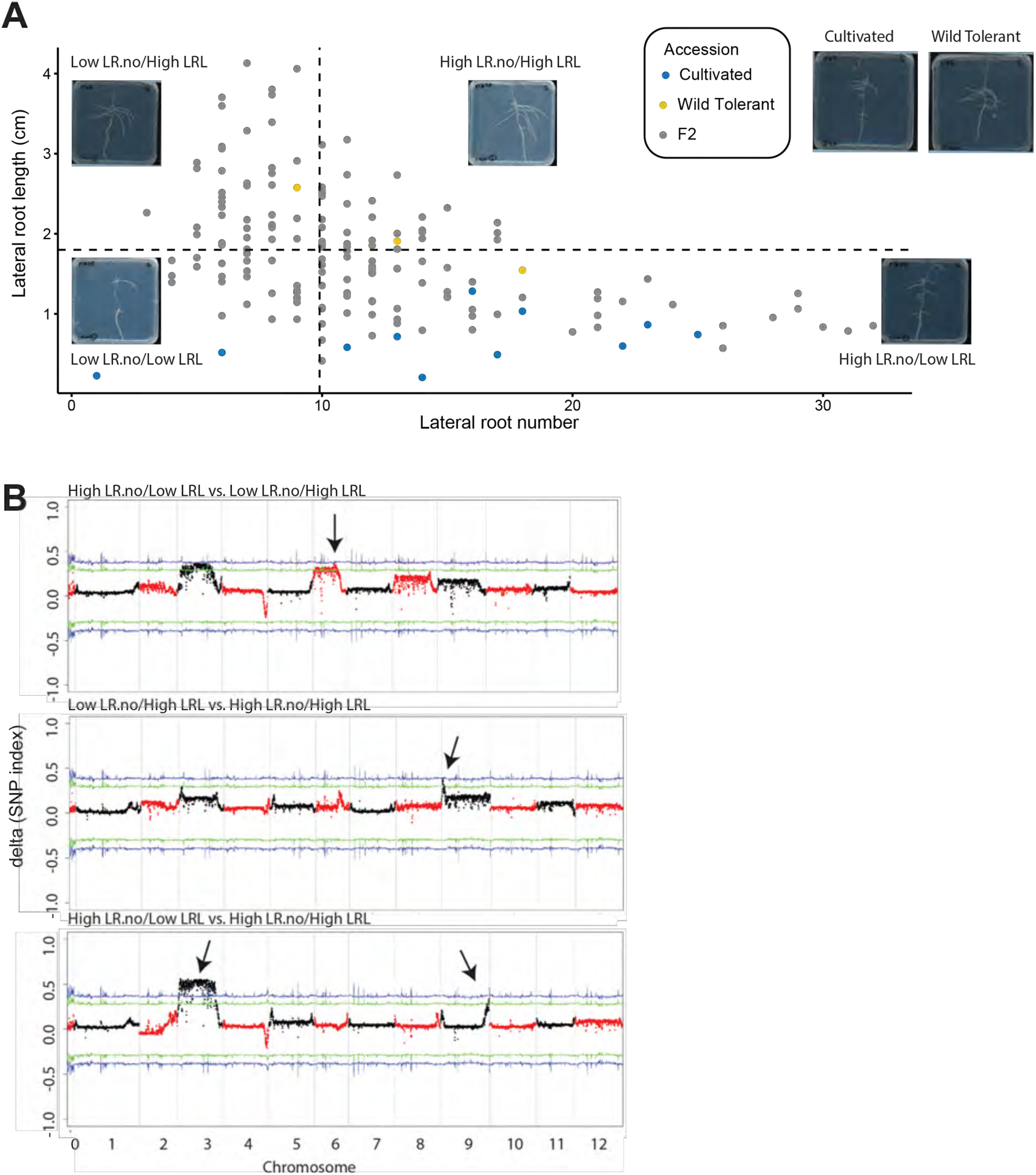
Segregating F2 populations displays four distinct phenotypic bulks for root system architecture. **(A)** Graph showing root system architecture of 180 F2 individuals alongside their parents under salt stress for lateral root length and lateral root number. The dashed lines used to show the threshold criteria in selecting different bulks, which was 1.8 cm for lateral root length and 10 for the lateral root number. Using these criteria, we identified four distinct phenotypic bulks, as indicated in the figure with representative images provided for each bulk and their parents. **(B)** Representative bulks comparison with most significant changes are shown. Distribution of Δ (SNP index) between the two bulks are shown across the *S. pimpinellifolium* genome. The statistical confidence intervals under the null hypothesis of no QTLs were indicated as green (*p* < 0.05) and blue (*p* < 0.01) lines.

We performed multiple bulk comparisons to identify genomic regions showing the most variation (**Fig. 4B and Fig. S8**). We found significant associations on chromosomes 3 and 6 with lateral root elongation (high LR.no/low LRL vs. low LR.no/high LRL, **Fig. 4B**) and chromosome 9 with lateral root emergence (low LR.no/high LRL vs. high LR.no/high LRL, **Fig. 4B**). Similarly, we observed a region towards the end of chromosome 9 and chromosome 3, associated with overall lateral root development under salt stress (high LR.no/low LRL vs. high LR.no/high LRL, **Fig. 4B**). These results suggest that the genes positioned on chromosomes 3, 6 and 9 are contributing to the differences in lateral root development between wild tolerant and cultivated tomato.

To narrow down the selection of candidate genes within the identified genetic regions, we overlaid the results from BSA and GWAS. We identified nine overlapping loci, and 22 gene coding regions that are putatively associated with lateral root development under salt stress (**Table 1, Table S3**). The candidate genes included: two ABC transporter family member genes (*Spim03g102820* and *Spim05g145990*), associated with the maintenance of main root length and total root system (MRLpTRS.D1&D2. SALT and TRS.D4.SALT.01); two N-acetylglucosaminyltransferase family genes (*Spim03g108680* and *Spim05g145980*) associated with the average lateral root length and total root system (aLRL.D1.SALT and TRS.D4.SALT); transcription factor bHLH87-like (*Spim03g110510*) and AP2-like ethylene-responsive transcription factor TOE3 (*Spim09g252830*), associated with the maintenance of average lateral root length and main root length per total root system (aLRL.D1.SALT and MRLpTRS.D3&D4.SALT). Additionally, we identified genes involved in the regulation of cellular processes such as those encoding a mannan synthase (*Spim09g253730*), a carboxypeptidase (*Spim09g253210*), a subtilisin-like protease (*Spim06g186120*), and a 3’-5’ exonuclease domain-containing protein (*Spim03g110520*), associated with maintenance of main root length and average lateral root length per total root system (MRLpTRS.D4.SALT, aLRLpTRS.D2.SALT, and aLRL.D1.SALT). The L-ascorbate peroxidase gene (*Spim09g252840*) was associated with maintaining the main root length per total root system (MRLpTRS.D3&D4.SALT). Additionally, we identified genes encoding two kinases and three proteins of unknown function, including one containing a domain of unknown function (DUF1677, *Spim06g186130*), which were associated with maintaining either lateral root or main root length per total root system (aLRLpTRS.D2.SALT). Although variation within the natural diversity panel is not guaranteed to be captured within bulk-segregant analysis due to differences in allele frequencies and causal genes, we identified overlapping genetic elements contributing to root development under salt stress between GWAS and bulk-segregant analysis.

**Table 1.** List of 22 genes identified by cross-referencing GWAS and BSA analysis. If no genes were found within the chromosome regions indicated by the GWAS, we extended the search by 10 kb before and after the original SNP locations.

| Locus | 1st SNP | Last SNP | Gene ID | Chromosome | Start | End | Strand | Description |
| --- | --- | --- | --- | --- | --- | --- | --- | --- |
| MRLpTRS.D1&D2.SALT | 58311594 | 58318648 | Spim03g102820 | Chr03 | 58317465 | 58333842 | - | ABC transporter A family member 7-like |
| aLRL.D1.SALT | 62872139 | 62872729 | no candidate |  |  |  |  |  |
| +/-10k | 62862139 | 62882729 | Spim03g108680 | Chr03 | 62862616 | 62865458 | - | Core-2/I-branching beta-1,6-N-acetylglucosaminyltransferase family |
|  |  |  | Spim03g108690 | Chr03 | 62868995 | 62871907 | - | AP-3 complex subunit delta |
|  |  |  | Spim03g108700 | Chr03 | 62873276 | 62881576 | - | Histidine-tRNA ligase |
| aLRL.D1.SALT | 64342558 | 64343225 | no candidate |  |  |  |  |  |
| +/-10k | 64332558 | 64353225 | Spim03g110510 | Chr03 | 64340070 | 64341212 | - | Transcription factor bHLH87-like |
|  |  |  | Spim03g110520 | Chr03 | 64348724 | 64354128 | - | 3'-5' exonuclease domain-containing protein |
| TRS.D4.SALT | 2853323 | 2873370 | Spim05g145980 | Chr05 | 2855866 | 2859416 | - | Core-2/I-branching beta-1,6-N-acetylglucosaminyltransferase family |
|  |  |  | Spim05g145990 | Chr05 | 2864524 | 2873811 | + | ABC transporter G family member protein |
| aLRLpTRS.D2.SALT | 39476715 | 39476898 | no candidate |  |  |  |  |  |
| +/-10k | 39466715 | 39486898 | Spim06g186100 | Chr06 | 39466736 | 39467116 | - | Receptor-type tyrosine-protein like |
|  |  |  | Spim06g186110 | Chr06 | 39470274 | 39473093 | + | Ring-box protein 1a |
|  |  |  | Spim06g186120 | Chr06 | 39473899 | 39476229 | - | subtilisin-like protease |
|  |  |  | Spim06g186130 | Chr06 | 39481301 | 39481819 | - | Protein of unknown function (DUF1677) |
| MRLpTRS.D3&D4.SALT | 893689 | 899226 | no candidate |  |  |  |  |  |
| -10k/+10k | 883689 | 909226 | Spim09g252830 | Chr09 | 885469 | 888212 | + | AP2-like ethylene-responsive transcription factor TOE3 |
|  |  |  | Spim09g252840 | Chr09 | 893083 | 897835 | + | L-ascorbate peroxidase |
|  |  |  | Spim09g252850 | Chr09 | 901262 | 910459 | + | Kinase |
| MRLpTRS.D3&D4.SALT | 1235817 | 1235817 | no candidate |  |  |  |  |  |
| -10k/+10k |  |  | Spim09g253200 | Chr09 | 1239184 | 1244087 | + | DnaJ protein like |
|  |  |  | Spim09g253210 | Chr09 | 1244798 | 1246201 | + | Carboxypeptidase |
| MRLpTRS.D3&D4.SALT | 1278892 | 1308283 | Spim09g253260 | Chr09 | 1289015 | 1294003 | + | TIR domain-containing protein |
|  |  |  | Spim09g253270 | Chr09 | 1304169 | 1306382 | + | Protein kinase domain-containing protein |
| MRLpTRS.D4.SALT | 1688182 | 1701564 | Spim09g253720 | Chr09 | 1691928 | 1693028 | + | Unknown protein |
|  |  |  | Spim09g253730 | Chr09 | 1694719 | 1698430 | - | Mannan synthase |
|  |  |  | Spim09g253740 | Chr09 | 1699511 | 1699663 | + | Unknown protein |

### Identification of transcriptional patterns underlying differences in lateral root development under salt stress

To identify key time points at which wild tolerant and cultivated tomatoes differ in their lateral root development in response to salt stress, we imaged the development of their root architecture under control and salt stress conditions at 60-minute intervals using a SPIRO setup (Ohlsson et al. 2021)(**Fig. S9**). We observed that salt stress exposure immediately reduces lateral root length in cultivated tomatoes, while no significant differences in individual lateral root length were observed in wild tolerant (**Fig. S9A**). Significant decrease in individual lateral root growth rate was observed from 1,200 minutes after stress exposure onwards in cultivated tomatoes (**Fig. S9B**). On the other hand, there was no obvious difference in lateral root growth rate of wild tolerant until 5,000 minutes after stress exposure, after which salt exposed lateral roots showed an increased growth rate (**Fig. S9B**). Cultivated tomatoes developed new lateral roots throughout the experiment under both control and salt stress conditions, albeit at a reduced rate, thereby reducing the average emergence time from 3519 to 4488 minutes after stress exposure (**Fig. S9C**). Although the average emergence time in wild tolerant showed even smaller difference between control and salt stress conditions (2841 and 3173 minutes after stress exposure respectively), we did observe a clear delay in lateral root emergence of wild tolerant under salt stress compared to the control condition, with no lateral roots emerging until 2,000 minutes after stress exposure (**Fig. S9C**).

Guided by the time points at which we observed significant differences in lateral root growth and emergence, we performed RNA-Seq analysis of the transcriptional profiles of both cultivated tomato and wild tolerant roots collected at 500, 1000, 2000, 3000, and 4000 minutes after salt stress exposure (**Fig. 5, Table S4**). We identified 3,792 differentially expressed genes (DEGs) in wild tolerant (2,261 upregulated and 1,531 downregulated) and 4,032 DEGs in cultivated tomato (2,307 upregulated and 1,725 downregulated) (**Fig. 5A**). Among these changes, we found 1,467 genes as being upregulated in both accessions while 1,009 genes downregulated. To identify genotype-specific DEGs at specific time points, we compared DEGs across individual time points (**Fig. 5B**). WT showed highest DEGs at 500 min after salt stress exposure (1,796 genes with fold change ≥ 2-fold and adjusted p-value > 0.05), followed by 4,000 min (1,631) (**Table S5**). On the other hand, cultivated tomato showed the highest number of DEGs at 4000 min (2,018 genes with fold change ≥ 2-fold and adjusted p-value > 0.05), followed by 500 (1,989). While both accessions showed the most changes in transcript abundances during early and late time exposure to salt stress, both had the least changes at 3,000 min after salt exposure. To compare the overlap between time points, we generated UpSet plots (**Fig. 5C, Fig. S10**). This analysis revealed that most upregulated DEGs in each accession were detected at 4,000 minutes, while most downregulated DEGs occurred at 500 minutes (**Fig. 5C**). Additionally, we observed that the majority of DEGs in each accession are unique to one or two time points, with only a few DEGs commonly expressed across all time points. Using gene ontology (GO) enrichment analysis, we examined at which biological process they were involved in. Most DEGs at 500 minutes after salt exposure were enriched with genes involved in stress responses and metabolic processes in wild tolerant, or detoxification and transporter activities in cultivated tomato. Interestingly, at 4,000 minutes, the wild tolerant DEGs remained enriched with genes involved in stress responses and metabolic processes, similar to the early response to stress. In contrast, DEGs found in cultivated tomato showed significant enrichment with genes related to light response, regulatory functions in photosynthesis and cellular binding activities (**Fig. 5C, Table S6**).

**Figure 5.**
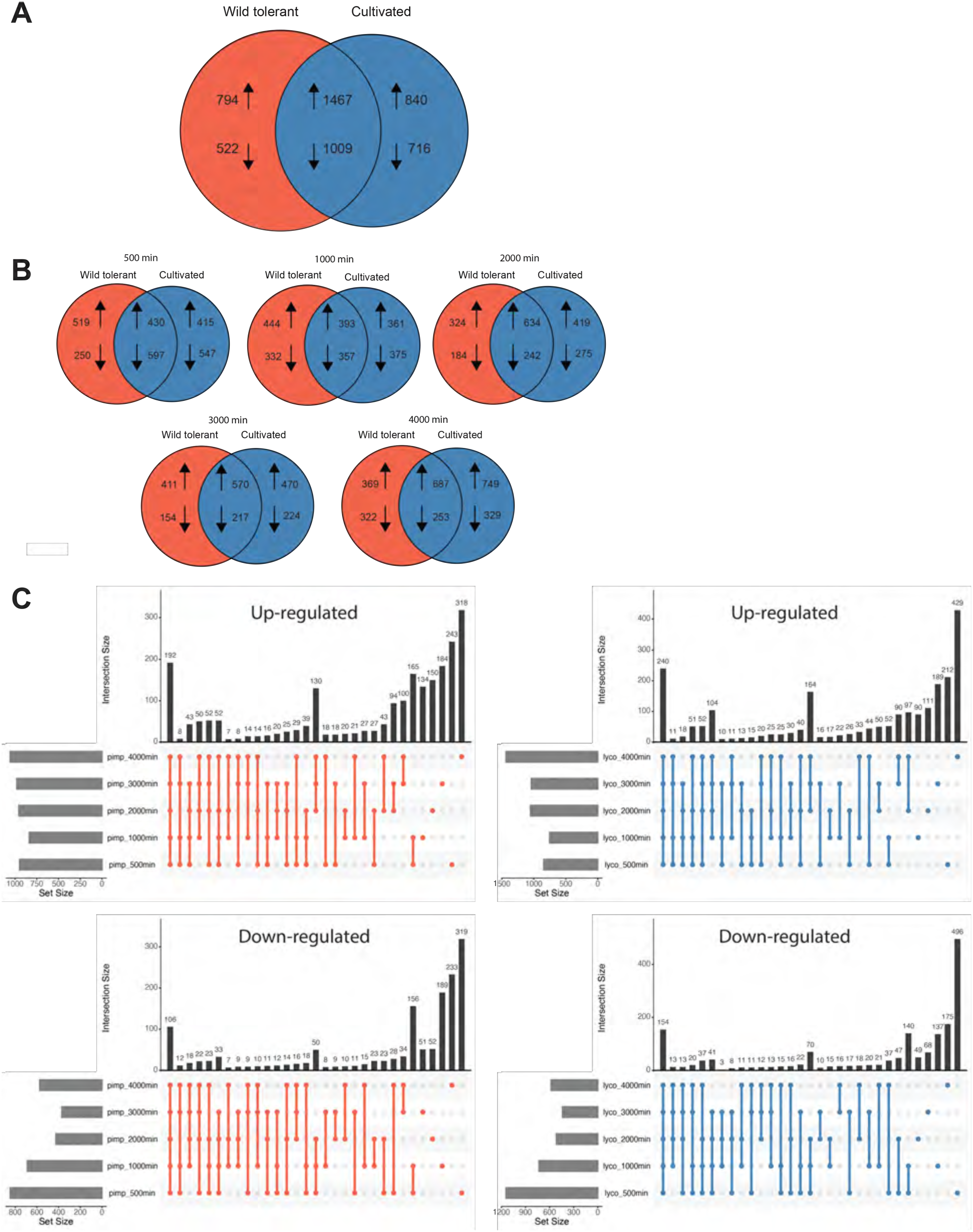
Differential expression analyses showing unique and common changes in transcriptome of the two tomato accessions in response to the salt stress. **(A)** Venn diagrams showing number of differentially expressed genes between the two accessions for all timepoints, and **(B)** across individual time points. **(C)** Intersection of numbers of DEGs under salt treatment relative to control condition across five timepoints. The graphs with orange dots represent DEGs in wild tolerant samples and graphs with blue dots represents cultivated tomato samples. The direction of arrow within each panel represents whether DEGs are significantly up- or down-regulated.

To uncover transcriptional patterns underlying lateral root differences between accessions, we conducted weighted gene correlation network analysis (WGCNA) (Langfelder and Horvath 2008) and identified 14 co-expression modules (**Fig. 6, Fig. S11**). We focused on modules containing GWAS/BSA candidates and those aligning with contrasting salt stress responses. The **yellow module** (2,697 genes) exhibited repression from 1,000 min post-stress, more pronounced in cultivated tomato (**Fig. 6A**). It included Spim05g145990, encoding an ABC transporter G family protein, emphasizing the role of cellular transport in root response. The **brown module** (2,907 genes) displayed initial repression in cultivated tomato, recovering over time, while wild tolerant showed a delayed but stronger resurgence (**Fig. 6B**). It contained Spim09g252850, a kinase-like gene linked to lateral root regulation. The **red module** (1,684 genes) showed early induction in cultivated tomato but later repression, while wild tolerant maintained upregulation (**Fig. 6C**). It included AP-3 complex subunit delta (Spim03g108690) and Ring-box protein 1a (Spim06g186110), involved in protein trafficking and interaction (Lee and Hwang 2014), (Abd-Hamid et al. 2020). The **turquoise module** (6,591 genes) was consistently more expressed in cultivated tomatoes, containing ten GWAS/BSA genes associated with protein homeostasis, signaling, and structural integrity (**Fig. 6D**). The **blue module** (3,427 genes) was upregulated in cultivated tomato under salt stress, with a transient upregulation in wild tolerant (**Fig. 6E**). It included genes related to stress response, oxidative protection, and protein modification. The black module (313 genes) was upregulated early in both accessions but showed no later treatment differences (**Fig. 6F**), featuring Spim03g102820, an ABC transporter A family member 7-like gene. These findings highlight that lateral root responses to salt stress depend on early stress-induced genes (e.g., yellow module) and constitutively differentially expressed genes (e.g., turquoise and blue modules).

**Figure 6.**
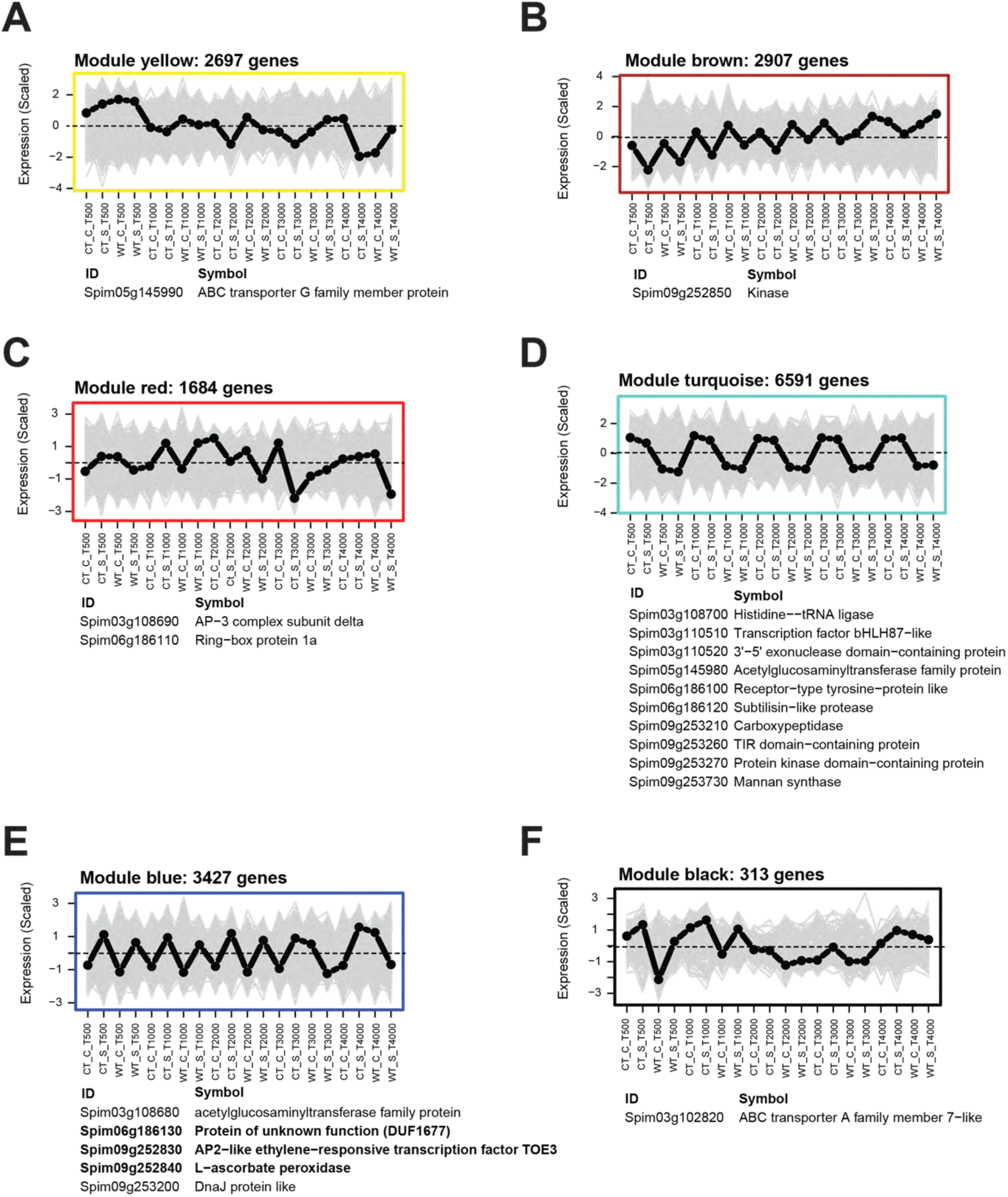
Co-expression network analysis showing distinct patterns among the two accessions across various developmental scales. WGCNA co-expression modules with scaled expression values (y axis) across different time points (x axis) **(A-F).** Black dotted line indicates eigengene expression profile. Gray line indicates expression values of all genes within the module. Most genes in these modules were positively correlated to the eigengene. Genes that are identified through GWAS and BSA (Table 1) and showing significant association with each module were listed below their respective modules, with bolded genes highlighting three of the four candidates identified by cross-referencing GWAS and BSA with RNA-Seq data. See Fig. S8 for a detailed analysis of all 14 modules.

### Identification of candidate genes highly associated with lateral root development under salt stress

To identify which of the 22 candidate genes (**Table 1**) exhibited transcriptional changes during root development, we cross-referenced them with the RNA-Seq data (**Table S4, Fig. S12**). While all were expressed across various time points, only four showed significant salt stress-induced changes in at least one accession (**Fig. 7A**): DUF1677 (Spim06g186130), AP2-like ethylene-responsive transcription factor TOE3 (Spim09g252830), L-ascorbate peroxidase (Spim09g252840), and an unknown protein (Spim09g253720). DUF1677 and TOE3 were transiently repressed in both cultivated tomato and wild tolerant, indicating their suppression may be critical for salt stress response. L-ascorbate peroxidase was consistently upregulated in cultivated tomato but only transiently in wild tolerant, suggesting its importance in lateral root development. The unknown protein was upregulated early in wild tolerant but not in cultivated tomato, highlighting a genotype-specific expression pattern.

**Figure 7.**
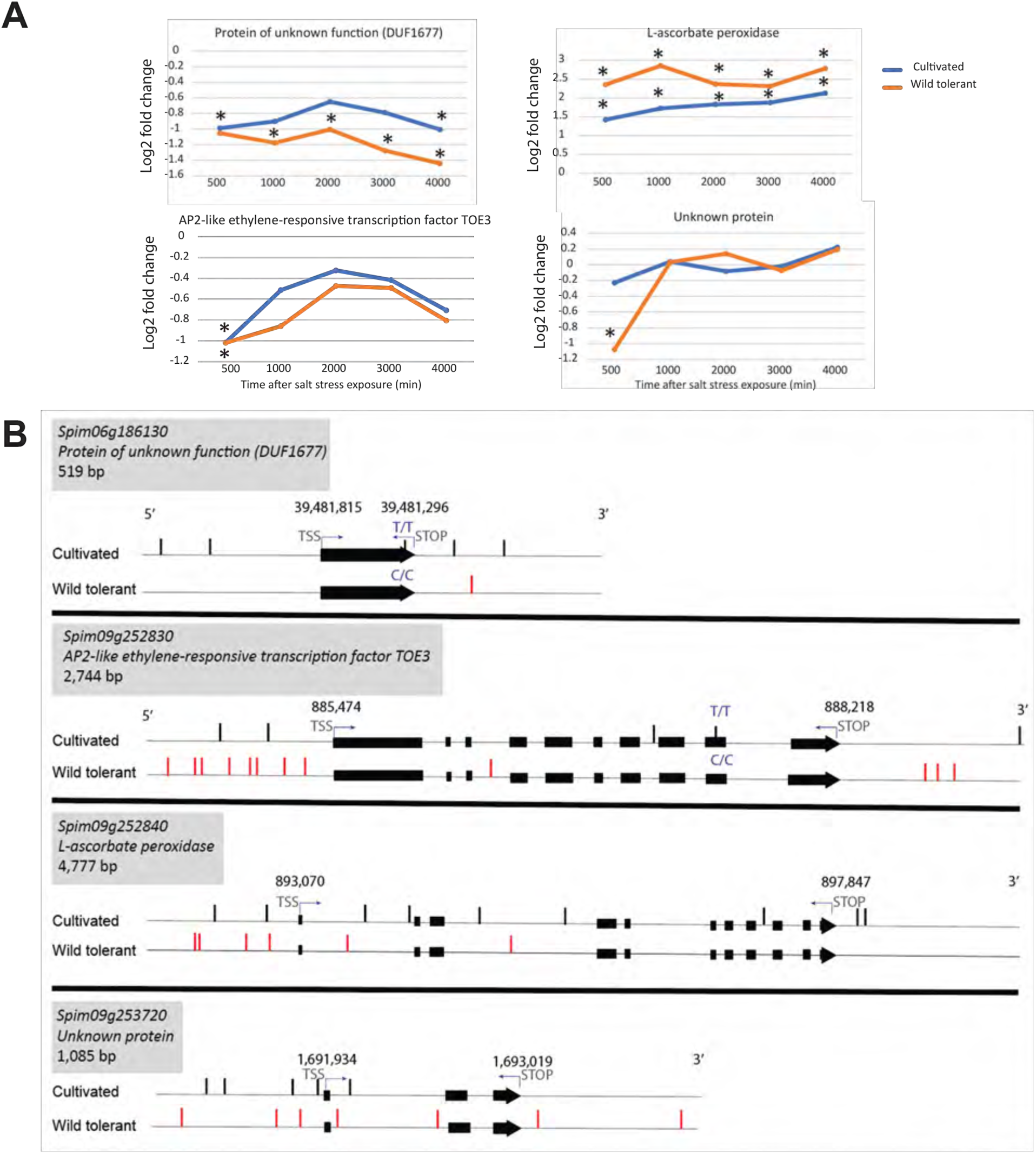
Allelic variants analysis revealed the two accessions are almost conserved in their coding sequence but not in the intronic region, as well as 5’ and 3’ UTRs. **(A)** Log2 fold changes are shown for the four genes identified by cross-referencing the three experiments, GWAS, BSA, and RNA-Seq, across all developmental time points for the two accessions. Stars indicate statistically significant changes in gene expression, defined as a fold change of ≥ 2 and an adjusted p-value of < 0.05. **(B)** Gene models are illustrated for the four genes, with unique SNP positions depicted as straight lines. The positions of the transcription start site (TSS) and STOP are indicated, with gene locations displayed above. SNPs resulting in missense mutations are depicted along with their corresponding variants in the other accession. See Table S3 for the comprehensive list of variant analyses conducted for the 22 genes identified from GWAS and BSA analyses (Table 1).

To assess allelic variation, we examined SNPs in these four genes using the Integrative Genomics Viewer (IGV) (**Fig. 7B**). SNPs were detected in their 5’ UTR, 3’ UTR, and intronic regions, suggesting roles in transcriptional regulation and mRNA stability. Two missense SNPs were identified: one in DUF1677 (Proline to Leucine) and one in TOE3 (Ala to Val or Leu to Phe), both differing from the reference *S. pimpinellifolium* LA2093 genome. Given the overall conservation of coding sequences between accessions, these findings suggest that regulatory variants likely drive differences in gene expression and function.

### ACC reduces root growth without affecting shoot Na⁺/K⁺ ratio

Given that TOE3, an AP2-like ethylene-responsive transcription factor, was identified as a key regulator of lateral root development under salt stress (**Fig. 7**), we hypothesized that ethylene is driving the different root architecture strategies in response to salt between cultivated and wild tomatoes. To test this, we analyzed root growth and Na⁺ accumulation in two tomato accessions treated with or without the ethylene precursor 1-aminocyclopropane-1-carboxylic acid (ACC). Across all accessions and conditions, ACC significantly reduced both main root and lateral root lengths (**Fig. 8, Fig. S13A-B**), while its effect on lateral root number varied. In cultivated tomatoes, low ACC concentrations significantly reduced lateral root number under both control and salt stress conditions, whereas high concentrations had no effect (**Fig. S13C**). In wild tomatoes, ACC had a non-significant effect under control conditions but significantly increased lateral root number under salt stress at both concentrations. To assess the physiological relevance of these changes, we calculated the STI for root traits (**Fig. 8B-D**). At low ACC concentrations, STI for main root growth significantly decreased in cultivated tomato but remained unchanged at high ACC concentrations (**Fig. 8B**). For lateral root growth, STI increased in cultivated tomato (non-significant at low ACC, significant at high ACC), while in wild tolerant, it significantly decreased (**Fig. 8C**). Lateral root number exhibited a non-significant STI increase in cultivated tomato and no clear trend in wild tolerant (**Fig. 8D**). These findings suggest that ethylene modulates root system architecture in response to salt stress, aligning with transcriptional changes observed in TOE3 (**Fig. 7**). To further explore ACC’s role in ion accumulation, we measured Na⁺, K⁺, and the Na⁺/K⁺ ratio using ICP-AES. ACC treatment resulted in a non-significant reduction of Na⁺ levels in roots and shoots (**Fig. S13D**) and a slight decrease in root K⁺ content, while shoot K⁺ levels remained unchanged (**Fig. S13E**). In cultivated tomato, the Na⁺/K⁺ ratio showed no change in roots at low ACC concentration but a non-significant increase at high ACC; in shoots, it decreased slightly at both concentrations (**Fig. 8E**). In wild tolerant, the Na⁺/K⁺ ratio showed a slight non-significant increase in roots and a mixed response in shoots. These results suggest that ACC-induced root architectural changes under salt stress influence Na⁺ and K⁺ accumulation, potentially contributing to stress resilience.

**Figure 8.**
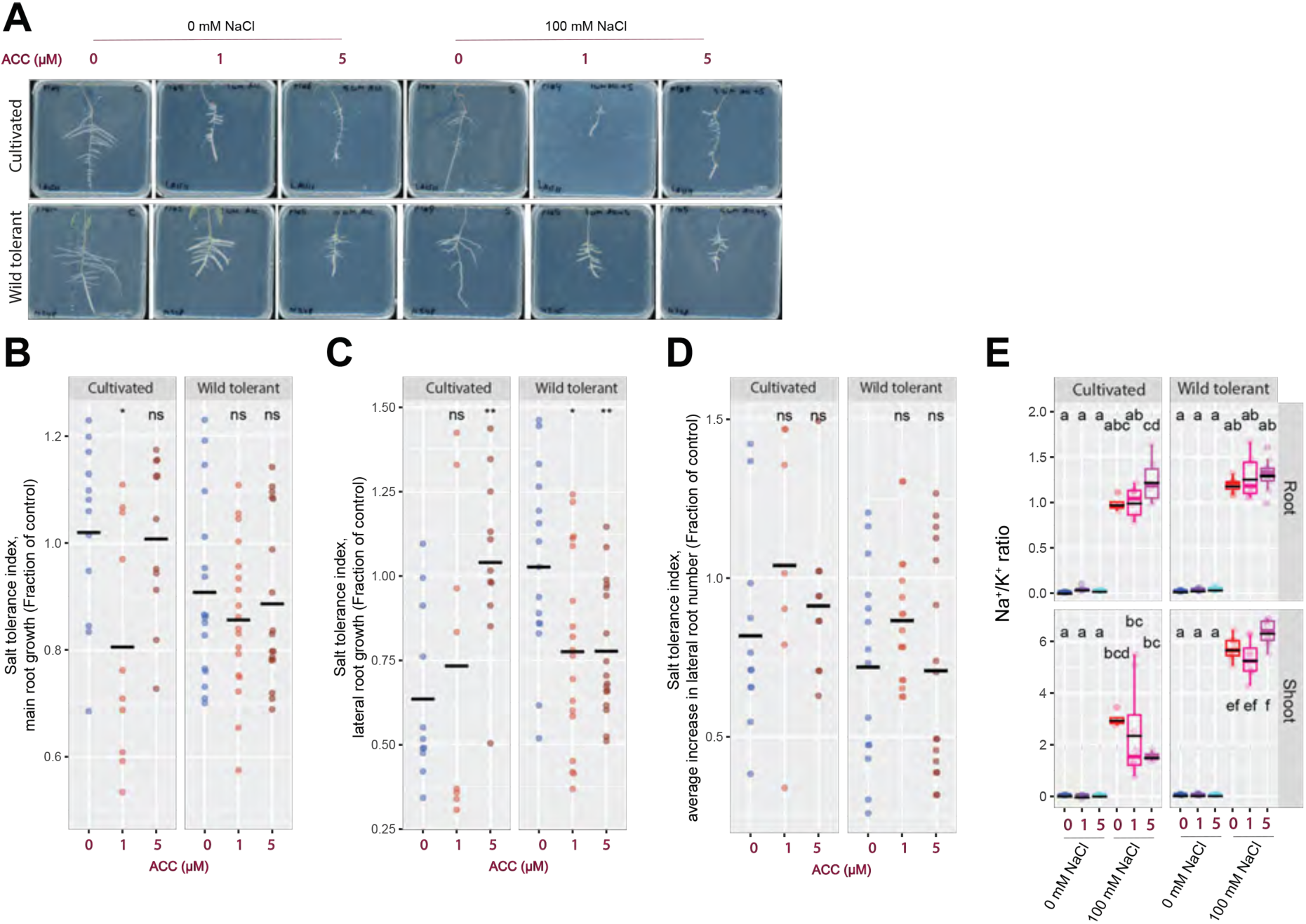
ACC treatment significantly reduces main root and lateral root length while lowering Na^+^/K^+^ ratio non-significantly in shoot of cultivated and wild tolerant tomato accessions. **(A)** Shown are representative images of 9-day-old tomato seedlings that experienced 5 days of treatment with 0 or 100 mM concentrations of NaCl supplemented with or without various concentrations of ACC as indicated in the figure. Salt tolerance indexes (STIs) are shown for the **(B)** main root, **(C)** average lateral root length, and **(D)** lateral root number. The STI was calculated by dividing the growth rate measured under salt stress by the growth rate measured under control condition for each accession. **(E)** Na^+^/K^+^ ratio of root and shoot of different accessions after 10 days on treatment plates. (B-E) Each dot represents individual replicas per accession and lines represent mean values in all. Statistical analysis was done by comparison of the means for all pairs using Tukey HSD test in all graphs. Levels not connected by the same letter are significantly different (P < 0.05).

### Ascorbate and H₂O₂ modulate root architecture and K⁺ retention under salt stress

Since APX was identified as a key candidate in lateral root development under salt stress (**Fig. 7**), we hypothesized that ROS homeostasis influences root system architecture under these conditions. To test this hypothesis, we analyzed root traits in two accessions under salt stress, with and without ascorbate (reducing agent) or H₂O₂ (oxidizing agent). Main root length and lateral root growth were largely unaffected by both treatments (**Fig. S14A-B**). However, in cultivated tomatoes, both ascorbate and H₂O₂ significantly increased lateral root number under control conditions but had no effect under salt stress, while wild tomatoes showed no significant response across treatments (**Fig. S14C**). Consistent with previous findings (**Fig. 2E-F**), cultivated tomato showed a higher STI for lateral root number, while wild tolerant had a higher STI for lateral root length (**Fig. 9A**, red highlights). Ascorbate had no significant effect on STI for main root length in cultivated tomato but caused a slight, non-significant decrease in wild tolerant (**Fig. 9A**, green font). STI for lateral root traits remained mostly unchanged, except for a slight, non-significant increase in lateral root length in wild tolerant. Similarly, H₂O₂ caused no significant changes in STI values, except for a slight increase in lateral root growth in cultivated tomato and a slight decrease in lateral root number (**Fig. 9A**). These results indicate that exogenous ascorbate and ROS influence root system architecture in a genotype-specific manner.

**Figure 9.**
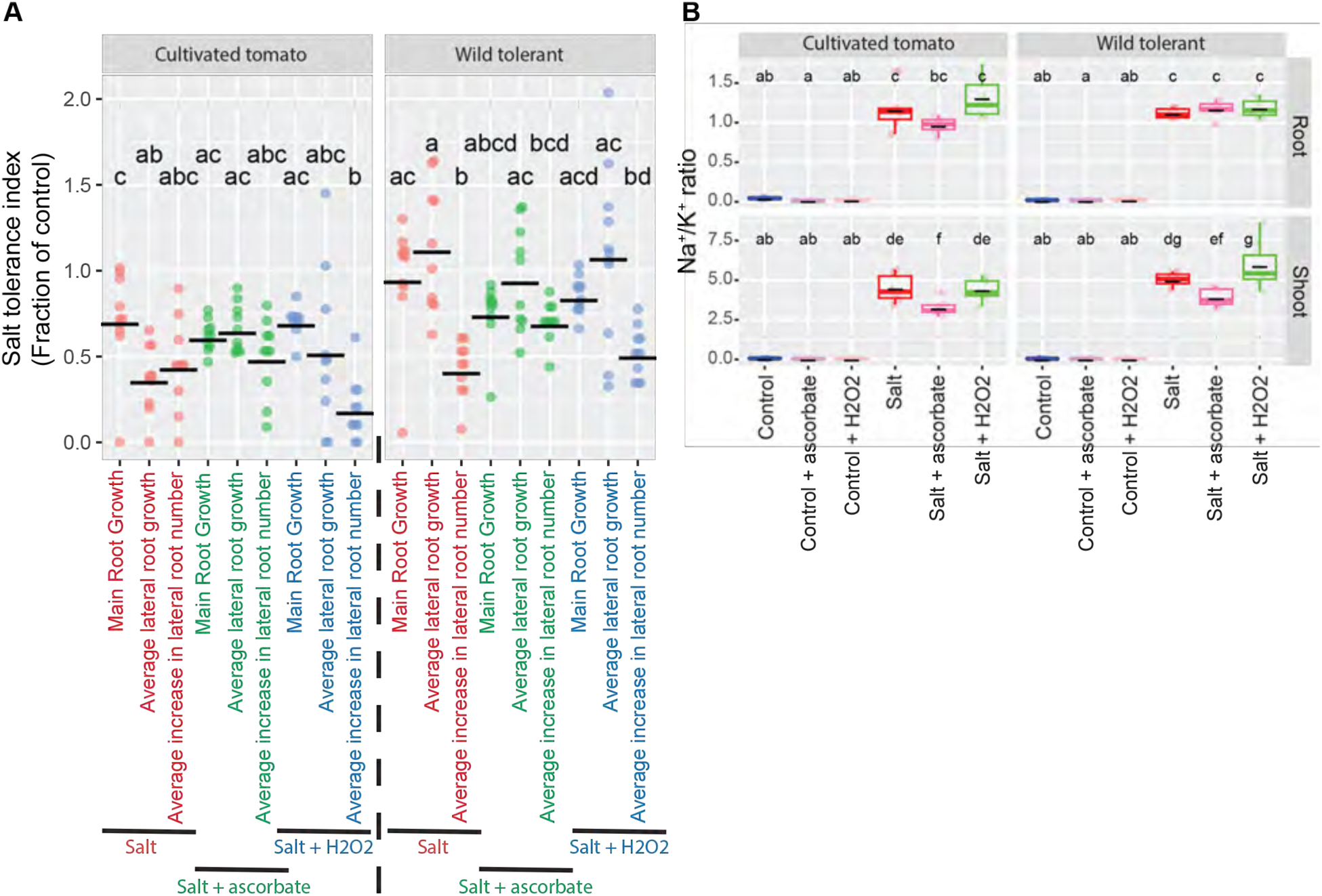
Ascorbate and H₂O₂ treatments show accession-dependent effects on root system architecture under salt stress, with ascorbate significantly reducing the shoot Na^+^/K^+^ ratio during salt stress. Analysis of root system architecture in tomato accessions treated with 0 or 100 mM NaCl, with or without the addition of 1 mM ascorbate or H₂O₂, as shown in the figure. **(A)** Salt tolerance indexes (STIs) are shown for the main root, average lateral root length, and lateral root number across the various treatments indicated in the figure. The STI was calculated by dividing the growth rate measured under salt stress by the growth rate measured under control condition for each accession. **(B)** Na^+^/K^+^ ratio of root and shoot of different accessions after 10 days on treatment plates. (A-B) Each dot represents individual replicas per accession and lines represent mean values in all. Statistical analysis was done by comparison of the means for all pairs using Tukey HSD test in all graphs. Levels not connected by the same letter are significantly different (P < 0.05).

To determine whether ascorbate and H₂O₂ treatments affect ion exclusion and Na⁺ and K⁺ homeostasis, we measured ion content in roots and shoots using ICP-AES (**Fig. S14D-E**). Neither treatment significantly altered Na⁺ content in either accession (**Fig. S14D**). Under control conditions, both treatments slightly increased root K⁺ content in cultivated tomato, while no significant changes were observed in wild tolerant. Under salt stress, ascorbate caused a slight but non-significant increase in root K⁺ content in both accessions, while H₂O₂ had no effect (**Fig. S14E**). In shoots, ascorbate led to a slight, non-significant K⁺ increase in cultivated tomato, whereas H₂O₂ had no detectable effect. In wild tolerant, ascorbate did not affect shoot K⁺ levels under control conditions, but H₂O₂ caused a minor increase. Under salt stress, shoot K⁺ content changes mirrored those in cultivated tomato (**Fig. S14E**). Regarding the Na⁺/K⁺ ratio, neither treatment significantly altered root ion balance, but ascorbate significantly reduced the Na⁺/K⁺ ratio in shoots under salt stress in both accessions (**Fig. 9B**). These results suggest that ascorbate enhances salt tolerance primarily by maintaining cellular K⁺ homeostasis.

### H₂O₂ accumulation under salt stress displays distinct responses among the two accessions

Given that we identified APX as a key candidate gene involved in lateral root development under salt stress (**Fig. 7**), and observed APX differential transcriptional responses among accessions in the transcriptome profiling, we hypothesized that the two accessions exhibit distinct H₂O₂ accumulation patterns upon salt stress in roots that might explain their distinct root architecture. To test this, we imaged H₂O₂ levels in the roots and shoots of both accessions over a time series under salt stress (**Fig. 10**). This analysis revealed that a spike of ROS accumulation occurs in the roots of the cultivated tomato at 20 and 30 minutes after salt exposure (p = 0.1), which then declined compared to corresponding mock treatments at 240 min (**Fig. 10A**). A similar pattern was not observed in the roots of the wild tolerant tomato (**Fig. 10A**). In contrast, the shoots of the cultivated tomato exhibited a peak in accumulation at 20 minutes (p = 0.1), while the shoots of the wild tolerant tomato showed a similar peak earlier at 10 minutes (p = 0.2), which continued to increase until 30 minutes after salt exposure (**Fig. 10B**). By 240 minutes post-exposure, both accessions displayed baseline levels of H₂O₂ in their shoots (**Fig. 10B**). Overall, these findings suggest that the two accessions exhibit distinct H₂O₂ signaling dynamics in roots in response to salt stress, which may contribute to their differences in root architecture.

**Figure 10.**
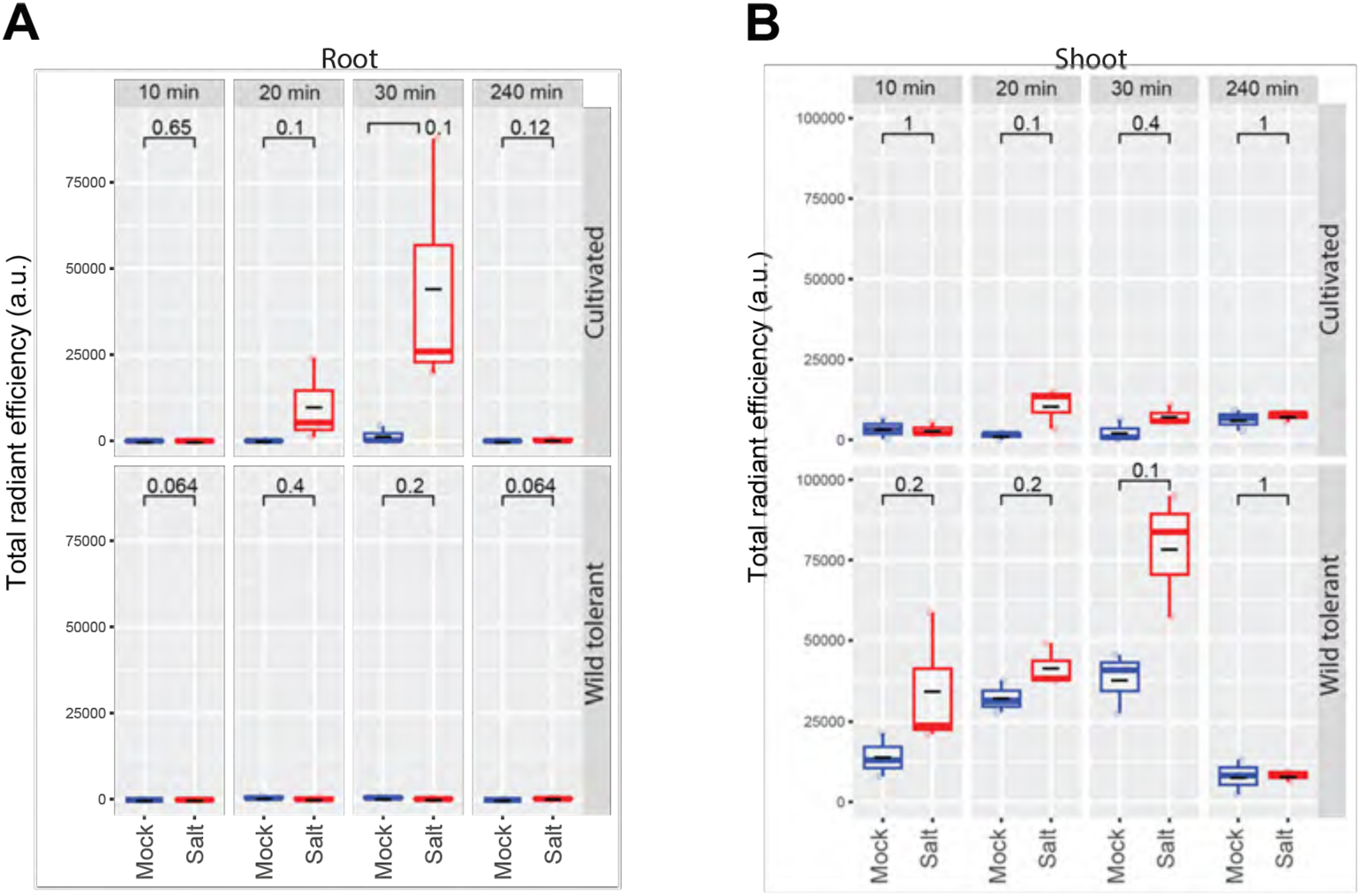
H_2_O_2_ profiling in roots shows distinct responses among the two accessions under salt stress. Quantification of H₂O₂ levels in the **(A)** roots and **(B)** shoots of two tomato accessions exposed to 0 or 100 mM NaCl at specific time points, as indicated in the figure. Salt stress was applied by fumigating the plates with a Peroxy Orange 1 (PO1) solution, a H₂O₂-specific dye, containing 100 mM NaCl, while mock-treated plates received only the PO1 solution. Imaging was performed using the IVIS Lumina S5 platform with Living Image 4.7.3 software in acquisition mode. The raw integrated density of regions of interest (ROIs) was quantified using ImageJ. Differences between mock and salt-treated conditions within each accession were analyzed using one-way ANOVA, with n = 3 per accession and condition. a.u. refers to arbitrary units.

## Discussion

Harnessing the inherent genetic diversity found in wild relatives, traditional landraces, and modern cultivars is essential for ensuring food security, a critical priority for crop enhancement initiatives. Genetic diversity within the wild progenitors of our cultivated crop species has been largely recognized as an important resource for increasing climate resilience of modern-day agriculture (Kovach and McCouch 2008; Morton et al. 2023). In tomato, GWAS has been used for detecting genetic variations underlying complex agronomic and morphological traits (Bui et al. 2015; Alaguero-Cordovilla et al. 2018; Rodriguez et al. 2020; Tripodi et al. 2021; Ye et al. 2021; Hakla et al. 2024). Although various components of salinity tolerance have been identified in S. pimpinellifolium (Rao et al. 2015; Wang et al. 2020b; Morton et al. 2023), this is the first study that explores root system architecture-derived mechanisms of salt tolerance.

We used a previously developed natural diversity panel of 220 diverse wild tomato accessions to identify alleles associated with root system responses to salt stress. A total of 191 significant associations corresponding to 55 genetic loci have been identified, of which nine showed overlap with BSA and RNA-seq studies. Moreover, we have identified accession LA2540 (wild tolerant) to be salt tolerant, which interestingly was identified as one of the more resilient *S. pimpinellifolium* accessions in field and greenhouse-based studies (Morton et al. 2023). Unlike in Arabidopsis, where natural variation in salt-induced changes in root architecture was observed for both main and lateral root based responses (Julkowska et al. 2017), we have identified that salt-induced changes in *S. pimpinellifolium* root architecture are mainly associated with lateral root development (**Fig. 1, Fig. S2**). We identified a cultivated tomato accession, LA1511, that showed complementary response in its lateral root development to the wild tolerant tomato (**Fig. 2**). Through generation of F1 hybrids between wild tolerant and cultivated tomato, we determined that the lateral root traits of both cultivated and wild tolerant accessions can be combined and act on complementary aspects of salinity tolerance. F1 hybrid exhibited an increase in STI for average lateral root length when compared to each parent (**Fig.3F**). Additionally, F1 maintained a lower Na^+^/K^+^ ratio, mimicking the cultivated parent and prioritizing tolerance by exclusion of sodium in the shoot (Fig. 6H) (Munns and Tester 2008). Together, these findings suggest that combining the complementary strategies of salinity tolerance, based on complementary characteristics of lateral root development, results in increased resilience of F1 hybrids. While previous studies have identified *S.pimpinellifolium* as a resource for salt tolerant traits (Rao et al. 2015; Razali et al. 2018; Martínez-Cuenca et al. 2020), we show that dissecting responses to salt stress through root architecture can result in identification of complementary traits that can be a source of further potentiation of salt stress resilience.

BSA identified three genomic regions linked to root growth maintenance under salt stress, and integration with GWAS revealed 22 candidate genes (**Table 1**). While only four showed significant transcript changes (**Fig. 7, Fig. S11**), 17 clustered in WGCNA co-expression modules (**Fig. 6**). Turquoise module contained genes involved in cellular homeostasis, including carboxypeptidase (Spim09g253210) and mannan synthase (Spim09g253730), key players in cell wall remodeling (Website; Voiniciuc 2022; Zhang et al. 2024), subtilisin-like protease (Spim06g186120), thus far mainly studied in the context of plant-pathogen interactions (Figueiredo et al. 2014), and 3’-5’ exonuclease domain-containing protein (Spim03g110520), implicated in DNA quality control (Shevelev and Hübscher 2002). Subtilases, known for their glycosylation-dependent activity, are critical for lateral root emergence, as seen in *Arabidopsis* where AIRS subtilase promotes lateral root outgrowth by degrading structural proteins in the extracellular matrix (Neuteboom et al. 1999). While mannan synthase remained repressed under stress, subtilisin-like protease exhibited late-stage upregulation (**Fig. S11**), suggesting distinct regulatory roles in lateral root adaptation to salt stress.

AP2-like ethylene-responsive transcription factor TOE3 (Spim09g252830) (**Fig. 6**) showed significant transcript changes under salt stress (**Fig. 7**) and co-localized within chromosome 9 GWAS/BSA regions (**Table 1**). Ethylene-responsive transcription factors regulate stress adaptation, often acting as molecular switches between stress tolerance and growth inhibition (Dubois et al. 2018). Ethylene’s role in salt tolerance varies—while enhancing resilience in *Arabidopsis*, it suppresses salt tolerance in *Oryza sativa*, where RNAi knockdown of OsEIL1/OsEIL2 improves survival (Cao et al., 2006; Achard et al. 2006; Jiang et al. 2013; Peng et al. 2014). Here, we observed that for both tomato accessions, ACC treatment significantly reduced main and lateral root growth (**Fig. 8, Fig. S13**) but had no significant impact on the shoot Na⁺/K⁺ ratio, suggesting ethylene limits root exposure to salt rather than directly affecting ion homeostasis. One possible mechanism is the growth restriction hypothesis, where ethylene reduces root surface area, minimizing Na⁺ uptake. TOE3 expression was initially repressed in response to salt stress before recovering at later time points (**Fig. 7**), indicating that lateral root development requires dynamic ethylene signaling. The early repression of TOE3 may be essential for lateral root initiation, followed by increased expression during adaptation. This aligns with findings in *Arabidopsis*, where ethylene results in early root growth inhibition through inhibiting cell proliferation and disturbance of microtubules in elongating cells (Street et al. 2015; Wang et al. 2018), but later facilitates root growth through promoting cell elongation via auxin-ethylene-ABA crosstalk (Růzicka et al. 2007; Thole et al. 2014). Together, these results support a model where ROS homeostasis, ethylene signaling, and cell wall remodeling coordinate lateral root development under salt stress. Future studies should explore post-transcriptional regulation of these candidate genes and how ethylene interacts with other hormonal pathways in salt adaptation.

Salinity stress leads to oxidative stress and cellular damage. The antioxidant system mitigates oxidative damage by counteracting the generation of reactive oxygen species (ROS) (Wang et al. 2013; Ahanger et al. 2017). Salt tolerance is positively correlated with the cell’s antioxidant capacity (Shalata and Neumann 2001; Munns and Tester 2008; Zhang et al. 2012; Wang et al. 2013; Dinneny 2019; Hasanuzzaman et al. 2019; Thabet et al. 2021), additionally supported here by an increase in the expression of ascorbate peroxidase throughout our experiment (**Fig. 7**). The constitutive expression of ascorbate peroxidase can be explained by two potential mechanisms. First, the initial reactive oxygen species (ROS) generated by salt stress trigger root cells to enhance their antioxidant response. Second, the emergence of lateral roots requires cell rupture, which generates another wave of ROS signal in a localized area of the root environment, further amplifying the increase in transcript levels. These potential mechanisms highlight the complex interplay between stress responses and root development. While ascorbate and H₂O₂ treatments showed accession-dependent effects on root system architecture under salt stress, we observed that ascorbate treatment significantly reduced the Na^+^/K^+^ ratio, primarily by aiding in K^+^ retention during salt stress (**Fig. 9, Fig. S14**). Previous studies have demonstrated that the Arabidopsis K^+^ efflux channel GATED OUTWARDLY-RECTIFYING K^+^ CHANNEL 1 (GORK1, AT5G37500) is activated by hydroxyl radicals (HO•) under salt stress and pathogen attack (Demidchik et al. 2010). Using electron paramagnetic resonance spectroscopy and electrophysiological techniques, it was shown that high NaCl concentrations (100 and 250 mM) stimulate HO• production, activating K^+^ channels and causing K^+^ efflux from roots. From our root transcriptomics data, we identified three outward-rectifying K^+^ channels that were differentially expressed in one or both tomato accessions (**Table S7**). Two of these channels (Spim03g099320 and Spim11g313920, both annotated as potassium channel SKOR) were downregulated under salt stress, while the third (Spim05g165150, annotated as potassium channel SKOR-like) was upregulated. Notably, Spim11g313920 shares the highest protein identity with Arabidopsis GORK1 (70%), followed by Spim05g165150 (68%), and Spim03g099320 (59%). Spim11g313920 is downregulated at most time points in our study in both accessions. GORK1 is known to contribute to guard cell membrane potential hyperpolarization by allowing K^+^ efflux, a process critical for resetting membrane potential during ion transport cycles. Under salt stress, GORK1 is expected to be upregulated to counterbalance Na^+^ influx at the cost of K^+^ efflux, maintaining cellular ionic balance. However, in our study, its downregulation suggests an adaptive response to retain K+ in the cytosol, preserving cellular integrity. Since both tomato accessions here demonstrate salt tolerance through distinct lateral root traits, we propose that the addition of ascorbate enhances the downregulation of K^+^ efflux channels, supporting K^+^ retention in the cytosol. This process helps maintain cellular integrity and promotes lateral root development under salt stress. However, further studies are needed to investigate how ascorbate influences the functionality of K^+^ efflux channels beyond changes in gene expression, which is outside the scope of this study. Although both accessions exhibited early H₂O₂ accumulation in shoots in response to salt, the distinct H₂O₂ signaling patterns observed in the roots of the two accessions suggest differential oxidative signaling dynamics that may regulate lateral root development under salt stress. The early and sustained H₂O₂ accumulation in cultivated tomato roots at 20 and 30 minutes (**Fig. 10A**) may contribute to salt-induced lateral root emergence. In contrast, the wild-tolerant accession, which exhibits minimal and delayed H₂O₂ accumulation at 240 minutes, may employ a different strategy to modulate redox signaling, potentially facilitating lateral root elongation. These findings suggest that temporal differences in H₂O₂ accumulation could underlie the contrasting root architectural responses observed between the accessions.

In conclusion, our integrative approach successfully identified new and previously known candidate genes involved in lateral root development under salt stress conditions. Our additional physiological experiments provide compelling initial validation that ethylene signaling and the antioxidant scavenging system are pivotal factors in regulating lateral root development under salt stress. Future full genetic validation of the identified genes in both wild and cultivated tomato background will provide more insight into genotype x environment interactions and genetic context of lateral root development. Nevertheless, data generated in this study will prove invaluable for future functional validation studies aimed at elucidating the precise roles of these genes during early root development in response to salt stress, contributing to the broader objective of enhancing plant productivity under saline conditions.

## Materials and Methods

### Agar-based plate experiments

Tomato seeds were sterilized for 10 min with 50% bleach and rinsed five times using autoclaved milli-Q water, and germinated on ¼ strength Murashige and Skoog (MS) medium containing 0.5% (w/v) sucrose, 0.1% (w/v) 4-morpholineethanesulfonic acid (MES), and 1% (w/v) agar, with pH adjusted to 5.8 with KOH. After 24 h of vernalization at 4 ^◦^C in the dark, the plates were placed in the Conviron growth chamber with the light intensity of 130–150 µmol x m^−2^ x s ^−1^ in a 20 h light / 4 h dark cycle at 25 ^◦^C day / 20 ^◦^C night and 60% humidity. 4 days after germination, the seedlings were transferred to 1/4 MS media with and without 100 mM NaCl, as indicated in the figures. Each plate contained one tomato seedling for root system architecture analysis. The plates were scanned using EPSON scanner every 5 consecutive days, starting from day 4 after germination. To analyze root system architectural traits from the scanned plate images, we used SmartRoot plugin (Lobet et al. 2011) in ImageJ to manually trace the root, and extract root-related features in the CSV format followed by data analysis in R. The root system architecture assays for the initial experiment to evaluate the root system architecture responses of natural diversity panel were conducted over six experimental batches, with four biological replicates per genotype per condition. The quantified data averaged per genotype and condition is listed in Table S1. The original image data, together with the traced files can be found at https://doi.org/10.5061/dryad.nk98sf826. The R-script used to curate and process the raw phenotypic data can be accessed at (https://rpubs.com/mjulkowska/BIGpimp_RSA_salt). The R-notebooks for the analysis of individual follow-up experiments can be accessed at <u>RPubs - 20211201_M248M058LA1511_RSA_analysis</u>, <u>RPubs - 20230502_RSA_F1_100mM_Salt</u>, <u>RPubs - 20230612_F1_FW_DW_ICP_Salt</u>, <u>RPubs - 20230605_RSA_analysis_F2_100mM_Salt_d9_all_batches_final</u>). The data analysis and visualization has been performed using ggplot2, ggpubr, corrplot, ggbeeswarm, ggridges, gapminder, RColorBrewer, ColorRamps, doBy, reshape2, cowplot, plotly and tidyverse libraries (Kassambara; Keitt; Wickham 2016, 2020; Wilke 2016; Wei and Simko 2017; Wickham et al. 2019; Chen 2020; Sievert 2020; Categorical Scatter (Violin Point) Plots [R package ggbeeswarm version 0.7.2] 2023, doBy: Groupwise Statistics, LSmeans, Linear Estimates, Utilities).

### Genome wide-association study

The genome wide-association study has been performed using the data from 220 *S. pimpinellifolium* accessions using the ASReml script as described in (Morton et al. 2023). Briefly, the SNP data has been curated and imputed for missing and heterozygous SNPs, and ran using three PCA components as estimations for population structure, and kinship matrix calculated using GAPIT (Wang and Zhang 2021). The code used for running the ASReml-based script can be accessed at https://github.com/mmjulkowska/Pimp_RSA_GWAS_salt/blob/main/20211011_RSA_GWAS_prep.R andhttps://github.com/mmjulkowska/Pimp_RSA_GWAS_salt/blob/main/20220101_ASReml_prep_for_pimp_RSA.R. The ASReml-method GWAS output files can be accessed at https://doi.org/10.5061/dryad.nk98sf826. Additionally, we have used the entire SNP dataset to run EMMA X method (Kang et al. 2010), using 3 PCA components as estimations for population structure, and kinship matrix calculated using the essential SNP set. The code used for running the EMMAX-based script can be accessed at https://github.com/mmjulkowska/Pimp_RSA_GWAS_salt/blob/main/20240208_EMMAX_GWAS%20pipeline.md. The EMMA-X GWAS output files can be accessed at https://doi.org/10.5061/dryad.nk98sf826. The data was subsequently investigated for all SNPs with association p-value above genome-wide threshold of −log10(p-value) > 5, as well as minor allele frequency and overlap of associations across individual traits and neighboring SNPs. The list of strongly associated SNPs can be found in **Table S2**.

### Elemental analysis

The elemental analysis was performed using Inductively Coupled Plasma Atomic Emission Spectroscopy (ICP-AES). For analysis of Na^+^ and K^+^ ions in roots and shoots of plate-grown seedlings, the plants were grown on 1/4 x MS plates as described above. After 10 days of exposure to 0 or 100 mM NaCl, root and shoot tissues were harvested and measured for fresh weight, rinsed in milli-Q water and collected into separate paper bags that were dried at 60 ^◦^C for 2-3 days. Subsequently, the dry weight was recorded. Samples were digested in double distilled HNO_3_, followed by the addition of 60/40 nitric/perchloric acid (v/v), continuing incubation at 150 °C. Samples were processed for ICP-AES analysis using a Thermo iCap 7000 ICP-AES after being diluted to 10 ml with deionized water. Ion content was calculated per dry weight for each sample, followed by data analysis in R.

### Generating the F1 hybrid and screening the F2 population for variation in later root length and number

The F1 hybrid population was created through reciprocal crosses between two tomato varieties: the cultivated tomato (*S. lycopersicum* LA1511) and the wild tolerant tomato (*S. pimpinellifolium* LA2540). The F1 hybrid underwent self-pollination to produce the F2 segregating population. From this population, 180 F2 tomato seeds were used for Bulk Segregant Analysis (BSA). The seeds underwent processing similar to that described for the agar-based plate experiments mentioned above for screening root system architecture. Following the root system architecture analysis, we identified four distinct bulks within the F2 population based on lateral root length and number. These bulks were then subjected to DNA extraction.

### DNA extraction for genome sequencing

From each bulk, we pooled 25 individuals that had the best characteristic of their designated pool for DNA extraction from roots. Extraction of the DNA was carried out using Wizard® Genomic DNA Purification Kit (Promega, catalogue # A1125) according to the manufacturer instructions. The integrity of DNA samples was checked using the electrophoresis gel and the concentration was determined by measuring the absorbance at 260 nm in a Nanodrop. 500 ng/µl of DNA from each pool and the two parents were sent on dry ice to the Novogene (Sacramento, CA, USA) for genome sequencing.

### Bulk Segregant Analysis (BSA)

Raw paired-end reads were processed to trim adaptor and low-quality sequences using Trimmomatic (version 0.39) (Bolger et al. 2014). The processed reads were aligned to the reference *Solanum pimpinellifolium* genome (Wang et al. 2020a)) using BWA-MEM (0.7.17-r1188) (Li 2013) with default parameters. Following read mapping, duplicated read pairs in each sample were marked using Picard (v2.26.6; http://broadinstitute.github.io/picard/), and SNPs were then called from the marked alignment files using GATK (v4.1.4.1) (McKenna et al. 2010). The called SNPs were filtered using GATK with parameters ‘QD < 2.0 || FS > 60.0 || MQ <40.0 || MQRankSum < −12.5 || ReadPosRankSum < −8.0’.

For BSA-Seq analysis, in each comparison of two bulks, SNP index of each SNP site was calculated for each bulk by dividing the number of reads harboring the allele of one parent with the total number of reads covering the site. Δ(SNP index) was then calculated by subtracting the SNP index in one bulk from the SNP index in the other bulk. An average Δ(SNP index) in each 200-kb sliding window with a step size of 100 kb across the *S. pimpinellifolium* genome was calculated. The 95% and 99% confidence intervals of the Δ (SNP index) under the null hypothesis of no QTLs were also calculated following the method described in (Takagi et al. 2013). QTLs were then identified as genomic regions with the Δ (SNP index) larger than the threshold (*p* < 0.05).

### Root sample collection for RNA-Seq experiment

To determine the time points corresponding to both pre- and post-lateral root emergence in the presence and absence of 100 mM salt, we utilized the Smart Plate Imaging Robot (SPIRO) platform to capture images of tomato seedlings cultivated on agar plates at 30-minute intervals over the course of one week. Subsequently, we analyzed these images using SmartRoot, and processed in an R-pipeline that can be accessed at (https://rpubs.com/mjulkowska/tomato_LR_LA1511vsM248) to identify the time points at which lateral roots emerged. Based on this analysis, we selected five time points spanning before, during, and after lateral root emergence under both control and salt stress conditions, specifically at 500, 1000, 2000, 3000, and 4000 minutes after transfer to the treatment plates. Following this selection, we collected root samples, excluding the 0.5 cm root tip, from the two tomato accessions at the designated time points in three independent biological replicates. These samples were promptly frozen in liquid nitrogen and stored at −80°C for subsequent analysis.

### RNA isolation, library preparation, and sequencing

Total RNA from roots were extracted using RNeasy Plant Mini Kit (Qiagen, Invitrogen, catalog # 74904), including a cleaning step using RNase-free DNase to eliminate genomic DNA contamination. The integrity and quality of RNA samples were evaluated using the Qubit RNA IQ Assay Kit (ThermoFisher Scientific, catalog # Q33221) and its concentration were evaluated using Qubit RNA HS Assay Kit (ThermoFisher Scientific, catalog # Q32852). 500 ng/µl of total RNA from each sample was sent on dry ice to the Novogene (Sacramento, CA, USA) for library preparation using NEBNext Ultra™ II RNA Kit for Illumina (Catalog# E7770L) paired end x 150 bp unstranded sequencing using NovaSeq. RNA-seq data were processed, in brief, using “trimmomatic” (v0.32, parameter: LEADING:3 TRAILING:3 MINLEN:36, TruSeq3-PE.fa:2:30:10:2, (Bolger et al. 2014)) to remove adaptors and low-quality reads. Then, clean reads derived from *S. pimpinellifolium* and *S. lycopersicum* were mapped to the *S. pimpinellifolium* reference genome by Hisat2 (v2.1.0, parameter: --rna-strandness RF -p 60 --no-softclip, (Kim et al. 2019)). Afterwards, the BAM files were processed by FeatureCounts to quantify transcript abundance at the gene level (v2.0.1 (Liao et al. 2014), parameter: -s 1 -p -t mRNA -g ID -O). DESeq2 (v3.19, (Love et al. 2013)) was used to normalize read counts among the 60 samples and identify differentially expressed genes (DEGs). Here, we included the four timepoint (500min, 1000min, 2000min, 3000min, 4000min), two species (*S. pimpinellifolium* and *S. lycopersicum*), and treatments (Control and Salt) as the experimental design factors. Then condition-wise DEGs per time points per species were identified using the cutoff of adjusted p-value < 0.05. Intersections of DEGs among time-points and species were characterized by the UpsetR plot function in R (Conway et al. 2017). The effect of timepoints, species, and treatment on transcriptome response was examined by principal component analysis (PCA) using in-house R-scripts (https://github.com/Leon-Yu0320/Tomato-salt-stress-transcriptome). Briefly, the PC values were calculated using the top 25% most variable genes based on standard deviation (SD) the PCA plot was visualized with ggplot2 (Wickham 2016).

### Go enrichment analysis

GO term enrichment analysis was performed to identify over-represented GO terms in the biological process category using Blast2GO (v3.1.11) (Conesa and Götz 2008). The Fisher’s exact test with the FDR correction for multiple testing was used to determine statistical significance (adjusted *P* value < 0.05).

### Co-expression network analysis

Co-expression network modules were built with the weighted gene correlation network analysis (WGCNA) R package version 1.72-1 (Langfelder and Horvath, 2008). Normalized counts were used as an input and 75% of the most variable genes were used for analysis. A scale-free network was created by applying a soft threshold power of 5. We created an unsigned network using the blockwiseModules function with following parameters: corType = bicor, max PO utliers = 0.05, merge Cut Height = 0.4 and max Block Size = 25 000. Gene Ontology enrichment analysis was carried out with GoStats R package version 2.62.0 with a standard hypergeometric test, to evaluate the enrichment of categories in each module compared to the set of all genes used to create the co-expression network.

### Ethylene, ascorbate, and H_2_O_2_ treatments for root assay

To analyze root system architecture with ethylene treatment, we utilized the ethylene precursor 1-aminocyclopropane-1-carboxylic acid (ACC). ACC was incorporated into the ¼ MS agar plates at concentrations of 1 µM or 5 µM. After autoclaving the media, ACC was added to the cooled medium after filter sterilization. For ascorbate and H_2_O_2_ treatments, a concentration of 1 mM was used in the ¼ MS agar plates. To prevent ascorbate degradation, freshly prepared ascorbate solution was sprayed on the seedlings daily throughout the experiment. Similar to ACC, H_2_O_2_ was added to the cooled ¼ MS medium after filter sterilization.

### H_2_O_2_ imaging

The seeds were germinated for four days on ¼ strength Murashige and Skoog (MS) medium as described in the “Agar-based plate experiments” section above. Salt stress was applied by fumigating the plates with Peroxy Orange 1 (PO1; H_2_O_2_-specific dye; excitation/emission 540 nm/565 nm; Tocris Bioscience cat-1199576-10-7, Bristol, United Kingdom) solution containing 100 mM NaCl while mock plates were fumigated with only PO1 solution. 10 mL of 100 μM PO1 solutions were made in 10 mM phosphate buffer (pH7.4) containing 0.002% Silwet L-77 (LEHLE seeds, Round Rock, TX, USA). Plates were fumigated with fine mist of dye solutions at designated time points (10, 20, 30, 240 min after salt exposure) in a glass chamber (24.5 x 12.5 x 16.5) as described in (Fichman et al. 2019, 2022; Fichman and Mittler 2020, 2021). The imaging was done using the IVIS Lumina S5 platform via Living Image 4.7.3 software in acquisition mode (PerkinElmer, Waltham, MA, USA) (Fichman et al. 2019; Fichman and Mittler 2020, 2021; Myers et al. 2023) and the raw integrated density of regions of interest (ROIs) was calculated using Image J.

## Data availability

Raw genome sequencing reads from Bulk Segregant Analysis (BSA) and RNA-Seq raw reads have been deposited in the National Center for Biotechnology Information BioProject database under the accession numbers PRJNA1108418 and PRJNA1119579, respectively. The phenotypic data in both raw and processed state, and GWAS results can be accessed at https://doi.org/10.5061/dryad.nk98sf826.

## Acknowledgments

The authors would like to thank BTI and KAUST Greenhouse Teams for caring for the plants. The authors would like to thank Lidor Shaar-Moshe at University of Haifa for their constructive discussions that greatly enriched this work. The authors would like to acknowledge support from the NSF-IOS #2023310 (ADLN) and NSF-IOS #2102120 (ADLN), NSF Mathematical Biology #2244735 (MMJ). The majority of funding for this work was generously provided by KAUST, with baseline funding awarded to Mark Tester and BTI’s startup funds awarded to Magdalena Julkowska.

## Author contributions

M.R.I. and M.M.J. conceived the project and wrote the manuscript with inputs from all coauthors, M.M.J. performed GWAS, H.S. and M.M.J quantified the root architecture data for the GWAS population, M.R.I. conducted majority of the experiments including root system architecture analysis of selected accessions, generation of F1 hybrid and F2 segregating population, root system architecture analysis of F2 population for bulk segregant analysis, collected samples and extracted DNA for bulk segregant analysis, collected samples and analyzed ion content data from ICP-AES, performed root system architecture analysis for ethylene, ascorbate, and H_2_O_2_ treatments as well as conducted ICP-AES analysis, M.R.I. and M.M.J. both collected samples for the time-series RNA-Seq experiment, M.R.I. extracted RNA for time-series RNA-Seq experiment, E.C. and M.P. helped in ICP-AES, L.Y. and A.D.L.N. performed RNA-Seq DEGs analysis, D.K. performed Co-expression network analysis, J.Z. and Z.F. performed Bulk Segregant analysis and GO enrichment, D.M. and R.M. performed H_2_O_2_ imaging and data analysis, M.A.T. helped with providing infrastructure for the initial screen and initial funding for the experiments.

## Supplemental figures and tables legends

**Figure S1. The natural variation in Root System Architecture components in response to salt stress.** The 220 genotypes of *S. pimpinellifolium* were germinated on ½ MS agar plates for 4 days and subsequently transferred to media containing 0 or 100 mM NaCl (Control and Salt respectively). The seedlings were imaged every 24 h after transfer for 7 consecutive days. The Root Architecture of seedlings was quantified for the first 4 days after transfer, and inspected for changes throughout time for **(A)** Total Root Size, **(B)** Main Root Length, **(C)** Lateral Root Length, **(D)** Lateral Root Number, **(E)** Center of Gravity (calculated as the position of MR with most Lateral Root Length), and **(F)** ratio of Main Root Length per Total Root Size. The individual lines represent the genotype-specific mean, whereas the dashed lines represent the population average. The *, **, *** and **** represent the significant differences between the treatment at individual time points as tested with two-way ANOVA with p-value < 0.05, 0.01, 0.001 and 0.0001, respectively.

**Figure S2. Salt-induced changes in root biomass distribution between main and lateral roots are associated with loci on Chromosomes 3 and 4.** Genome Wide Association Study was performed on 9M SNPs using standard EMMA-X method. Only SNPs with minor allele frequency > 0.05 were used for plotting the Manhattan plot for associations identified with **(A)** ratio of main root length to total root size at 1 day and **(B)** two days post transfer to salt stress, with pseudo-heritability values of 0.116 and 0.195 respectively. The red line indicates the Bonferroni threshold, whereas the blue line indicates genome-wide suggestive threshold. **(C)** The significantly associated loci were inspected in further detail for the gene coding sequences within the regions with significantly associated SNPs. The region on chromosome 3 contained 29 SNPs with −log10(p-value) > 5. The three individual loci **(D-F)** identified on chromosome 4 contained 240, 39 and 85 SNPs with −log10(p-value) > 5. The association graph below each locus represents the association strength with the ratio of main root length to total root size at 1 day after salt stress imposition.

**Figure S3. Salt-induced changes in lateral root development are associated with loci on chromosomes 1, 3 and 10.** Genome Wide Association Study was performed on 9M SNPs using standard EMMA-X method. Only SNPs with minor allele frequency > 0.05 were used for plotting the Manhattan plot for associations identified with **(A)** average lateral root length and **(B)** lateral root number at one day post transfer to salt stress, with pseudo-heritability values of 0.145 and 0.358 respectively. The red line indicates the Bonferroni threshold, whereas the blue line indicates genome-wide suggestive threshold. **(C)** The significantly associated loci were inspected in further detail for the gene coding sequences within the regions with significantly associated SNPs. The region on chromosome 1 contained 27 SNPs, **(D)** the region on chromosome 3 7 SNPs, and **(E)** region on chromosome 11 contained 60 SNPs with −log10(p-value) > 5. The association graph below each locus represents the association strength with the average lateral root length at 1 day after salt stress imposition.

**Figure S4. Emergence of lateral roots and main root growth rate is associated with loci on chromosomes 12 and 22.** Genome Wide Association Study was performed on 9M SNPs using standard EMMA-X method. Only SNPs with minor allele frequency > 0.05 were used for plotting the Manhattan plot for associations identified with **(A)** lateral root emergence and **(B)** main root growth rate under salt stress, with pseudo-heritability values of 0.855 and 0.628 respectively. The red line indicates the Bonferroni threshold, whereas the blue line indicates genome-wide suggestive threshold. **(C)** The significantly associated loci were inspected in further detail for the gene coding sequences within the regions with significantly associated SNPs. The region on chromosome 12 contained 16 SNPs, whereas **(D)** the region on chromosome 11 contained 15 SNPs −log10(p-value) > 5. The association graph below each locus represents the association strength with the trait initially associated with that locus.

**Figure S5. High-confidence loci identified through GWAS.** Genome Wide Association Study was performed on 9M SNPs using the standard EMMA-X method. Only SNPs with minor allele frequency > 0.05 were used for plotting the Manhattan plot for associations identified with **(A)** lateral root number at 2 days after transfer to control conditions. The significantly associated loci on chromosome 5, consisting of **(B)** 15 and **(C)** 11 SNPs were inspected in further detail. **(D)** Associations with the number of lateral roots were identified on chromosome 1, and **(E)** the locus consisting of 14 highly associated SNPs has been inspected in more detail. The red line on Manhattan plots indicates the Bonferroni threshold, whereas the blue line indicates genome-wide suggestive threshold. The association graph below each locus represents the association strength with the trait initially associated with that locus.

**Figure S6. Wild tomatoes accumulate higher levels of Na^+^ in the shoot compared to cultivated tomato. (A)** Na^+^ and **(B)** K^+^ contents of root and shoot of different accessions after 10 days on treatment plates. Each dot represents an individual replicate per accession. Statistical analysis was done by comparison of the means for all pairs using Tukey HSD test. Levels not connected by the same letter are significantly different (P < 0.05).

**Figure S7. F1 individuals accumulate less Na^+^. (A-C)** Root system architecture of F1 individuals, their corresponding parents, i.e., cultivated and wild tolerant tomatoes along with wild sensitive tomatoes were investigated with and without 100 mM NaCl. **(D-E)** Na^+^ and K^+^ contents of root and shoot of different accessions after 10 days on treatment plates. Each dot represents individual replicas per accession. Statistical analysis was done by comparison of the means for all pairs using Tukey HSD test in all graphs. Levels not connected by the same letter are significantly different (P < 0.05).

**Figure S8. Remaining bulk comparisons.** Displayed here are the additional BSA results that were not shown in Figure 4.

**Figure S9. Wild tolerant and cultivated tomatoes show variation in lateral root elongation and emergence under the salt stress. (A)** Lateral root length, **(B)** lateral root length growth rate, and **(C)** lateral root number are shown for the two tomato accessions grown in agar plate under 100 mM NaCl. The plates were imaged using SPIRO setup at a 30-min interval for one week. Individual transparent lines indicate the growth trajectory of individual lateral roots, dashed line indicates the growth average per genotype per condition, whereas the shaded area indicates the standard deviation. The differences between Control and Salt conditions were tested within each genotype using one-way ANOVA, with *, ** and *** representing p-values below 0.05, 0.01 and 0.001, respectively. The growth dynamics of lateral roots were measured over two experimental replicates for 4 biological replicates per genotype per condition.

**Figure S10. Intersection of DEGs between WT and CT under Salt and Control conditions.** The upset plots highlight the intersection of numbers of DEGs between wild tolerant and cultivated tomato samples across five timepoints under salt and control conditions, respectively. The panel with dark blue dots represents samples under the control condition, while the panel with brown dots are Salt samples. The direction of arrows within each panel indicates whether the depicted genes are up- or down-regulated.

**Figure S11. Co-expression network analysis.** The rest of WGCNA co-expression modules are shown here, module 7 to 14.

**Figure S12. Gene expression profiles over time for genes identified by overlapping GWAS and BSA analyses in Table 1**. Log2 fold changes are shown for the 18 genes identified by cross-referencing the GWAS and BSA, across all developmental time points for the two accessions.

**Figure S13. ACC treatment significantly reduces both main root and lateral root lengths while causing a non-significant decrease in Na^+^ concentrations in the roots and shoots of both accessions under salt stress.** Root system architecture analysis of cultivated and wild tomatoes under 0 or 100 mM concentrations of NaCl supplemented with or without various concentrations of ACC, as indicated in the figure, are shown for main root growth rate **(A)**, average lateral root length **(B)** as well as average lateral root number **(C)**. Na^+^ **(D)** and K^+^ **(E)** content of root and shoot of different accessions after 10 days on treatment plates. (A-E) Each dot represents individual replicate per accession. (D-E) Lines represent mean values. The asterisks above the graphs in (A-C) indicate significant differences between control (i.e., 0 mM NaCl and 0 ACC) and other treatment by the Student ‘s t-test: *P < 0.05, **P < 0.01, ***P < 0.001, and ****P < 0.0001. Statistical analysis was done by comparison of the means for all pairs using Tukey–Kramer HSD test for (D-E). Levels not connected by the same letter are significantly different (P < 0.05).

**Figure S14. Ascorbate and H₂O₂ treatments exhibit accession-dependent effects on root system architecture under salt stress, with ascorbate contributing to K^+^ retention during salt stress.** Root system architecture analysis of cultivated and wild tomatoes under 0 or 100 mM concentrations of NaCl supplemented with or without ascorbate or H_2_O_2_, as indicated in the figure, are shown for main root growth rate **(A)**, average lateral root length **(B)** as well as average lateral root number **(C)**. Na^+^ **(D)** and K^+^ **(E)** content of root and shoot of different accessions after 10 days on treatment plates. (A-E) Each dot represents individual replicate per accession. (D-E) Lines represent mean values. Statistical analysis was done by comparison of the means for all pairs using Tukey–Kramer HSD test for (A-E). Levels not connected by the same letter are significantly different (P < 0.05).

**Table S1. Root System Architecture traits for *S. pimpinellifolium* diversity panel.** The 89 different phenotypes were recorded for 220 *S. pimpinellifolium* accessions. The traits presented in this table represent the average trait value as calculated based on 4 biological replicates. The trait abbreviations are as follow: aLRL - average Lateral Root Length; aLRLpTRS - ratio of average lateral root length to total root size; CoG - center of gravity calculated as the length of the main root that gives rise to most lateral root biomass; LRL - cumulative lateral root length; LRno - lateral root number, MRL - main root length; MRLpTRS - ratio of main root length to total root size; TRS - total root size. The abbreviations behind the 1st dot indicate either the time and treatment at which individual RSA parameters were recorded (0 - 4 - days post treatment application; C and S for Control and Salt stress respectively), or STI - stress tolerance index - calculated by dividing the trait value recorded under Salt stress conditions by value recorded under Control conditions. GR indicates the growth rate, estimated by fitting a linear curve into change of trait over time.

**Table S2. GWAS results.** All of the SNPs with −log10(p-value) > 5 ware collated into one table, from two GWAS methods used (ASReml - as used in (Awlia et al. 2021), and EMMA-X). The position of each SNP is indicated in Chr and Pos columns, corresponding to chromosome and position on the chromosome in base pairs as per S.pimpinellifolium genome annotation (Wang et al. 2020a). The distance between neighboring SNPs is indicated in the “dist” column, and distances below 10,000 bp are highlighted in yellow, while distances below 1,000 bp are highlighted in red. The Pval column indicates the P-value associated with each SNP, and MAF indicates the minor allele frequency of that particular SNP. The trait column indicates the specific trait association used, with aLRL representing average lateral root length, MRL main root length, LRL lateral root length, TRS total root size, perc - a percentage of main root length, CoG - center of gravity calculated for lateral root distribution along the main root length, Apical, Branched and Basal indicating the lengths of individual zones as absolute values or (perc) as percentage of the main root length. The traits recorded under Control conditions are colored in blue, while traits recorded under salt stress are colored with pink. The LOD is calculated as −log10(p-value). The Method represents whether an individual association was observed as significant using ASReml or EMMA-X method.

**Table S3. Analysis of allelic variants for the 22 genes that overlap between GWAS and BSA studies. Table S4. RNA-Seq raw reads and differentially expressed genes (DEGs).**

**Table S5. Differentially expressed genes (DEGs) for 500 and 4000 minutes.**

**Table S6. Go enrichment analysis for 500 and 4000 minutes.**

**Table S7. List of potassium channels and transporters with their gene expression profiles in tomato root transcriptomics.**

## References

Abd-Hamid N-A, Ahmad-Fauzi M-I, Zainal Z, and Ismail I. Diverse and dynamic roles of F-box proteins in plant biology. Planta. 2020:251(3):68.

Abrol IP, Yadav JSP, and Massoud FI. Salt-affected Soils and Their Management (Food & Agriculture Org.).

Achard P, Cheng H, De Grauwe L, Decat J, Schoutteten H, Moritz T, Van Der Straeten D, Peng J, and Harberd NP. Integration of plant responses to environmentally activated phytohormonal signals. Science. 2006:311(5757):91–94.

Ahanger MA, Tomar NS, Tittal M, Argal S, and Agarwal RM. Plant growth under water/salt stress: ROS production; antioxidants and significance of added potassium under such conditions. Physiol Mol Biol Plants. 2017:23(4):731–744.

Alaguero-Cordovilla A, Gran-Gómez FJ, Tormos-Moltó S, and Pérez-Pérez JM. Morphological Characterization of Root System Architecture in Diverse Tomato Genotypes during Early Growth. Int J Mol Sci. 2018:19(12). 10.3390/ijms19123888

Ashrafi H, Kinkade M, and Foolad MR. A new genetic linkage map of tomato based on a Solanum lycopersicum x S. pimpinellifolium RIL population displaying locations of candidate pathogen response genes. Genome. 2009:52(11):935–956.

Asins MJ, Bolarín MC, Pérez-Alfocea F, Estañ MT, Martínez-Andújar C, Albacete A, Villalta I, Bernet GP, Dodd IC, and Carbonell EA. Genetic analysis of physiological components of salt tolerance conferred by Solanum rootstocks. What is the rootstock doing for the scion? Theor Appl Genet. 2010:121(1):105–115.

Atkin RK, Barton GE, and Robinson DK. Effect of Root-growing Temperature on Growth Substances in Xylem Exudate of Zea mays. J Exp Bot. 1973:24(2):475–487.

Awlia M, Alshareef N, Saber N, Korte A, Oakey H, Panzarová K, Trtílek M, Negrão S, Tester M, and Julkowska MM. Genetic mapping of the early responses to salt stress in Arabidopsis thaliana. Plant J. 2021:107(2):544–563.

Bolger AM, Lohse M, and Usadel B. Trimmomatic: a flexible trimmer for Illumina sequence data. Bioinformatics. 2014:30(15):2114–2120.

Brozynska M, Furtado A, and Henry RJ. Genomics of crop wild relatives: expanding the gene pool for crop improvement. Plant Biotechnol J. 2016:14(4):1070–1085.

Bui HH, Serra V, and Pagès L. Root system development and architecture in various genotypes of the Solanaceae family. Botany. 2015:93(8):465–474.

Cao WH, Liu J, He XJ, Mu RL, Zhou HL, Chen SY, Zhang JS. Modulation of Ethylene Responses Affects Plant Salt-Stress Responses, Plant Physiology, Volume 143, Issue 2, February 2007, Pages 707–719, 10.1104/pp.106.094292

Categorical Scatter (Violin Point) Plots [R package ggbeeswarm version 0.7.2]. 2023.

Celik I, Gurbuz N, Uncu AT, Frary A, and Doganlar S. Genome-wide SNP discovery and QTL mapping for fruit quality traits in inbred backcross lines (IBLs) of solanum pimpinellifolium using genotyping by sequencing. BMC Genomics. 2017:18(1):1.

Chen B. Access Data from the USDA ARMS Data API [R package rarms version 1.0.0]. 2020.

Cione APP, Schmitt CC, Neumann MG, and Gessner F. The Effect of Added Salt on the Aggregation of Clay Particles. J Colloid Interface Sci. 2000:226(2):205–209.

Conesa A and Götz S. Blast2GO: A comprehensive suite for functional analysis in plant genomics. Int J Plant Genomics. 2008:2008:619832.

Conway JR, Lex A, and Gehlenborg N. UpSetR: an R package for the visualization of intersecting sets and their properties. Bioinformatics. 2017:33(18):2938–2940.

Demidchik V, Cuin TA, Svistunenko D, Smith SJ, Miller AJ, Shabala S, Sokolik A, and Yurin V. Arabidopsis root K+ efflux conductance activated by hydroxyl radicals: Single-channel properties, genetic basis and involvement in stress-induced cell death. J Cell Sci. 2010:123:1468–1479.

Dempewolf H, Eastwood RJ, Guarino L, Khoury CK, Müller JV, and Toll J. Adapting Agriculture to Climate Change: A Global Initiative to Collect, Conserve, and Use Crop Wild Relatives. Agroecology and Sustainable Food Systems. 2014:38(4):369–377.

Dinneny JR. Developmental Responses to Water and Salinity in Root Systems. Annu Rev Cell Dev Biol. 2019:35:239–257.

doBy: Groupwise Statistics, LSmeans, Linear Estimates, Utilities. Comprehensive R Archive Network (CRAN). https://cran.r-project.org/web/packages/doBy/index.html. Retrieved June 5, 2024

Dubois M, Van den Broeck L, and Inzé D. The pivotal role of ethylene in plant growth. Trends Plant Sci. 2018:23(4):311–323.

Ebert AW and Schafleitner R. Utilization of wild relatives in the breeding of tomato and other major vegetables. In. Crop Wild Relatives and Climate Change. (John Wiley & Sons, Inc: Hoboken, NJ, USA), pp. 141–172.

Fichman Y, Miller G, and Mittler R. Whole-plant live imaging of reactive oxygen species. Mol Plant. 2019:12(9):1203–1210.

Fichman Y and Mittler R. Noninvasive live ROS imaging of whole plants grown in soil. Trends Plant Sci. 2020:25(10):1052–1053.

Fichman Y and Mittler R. Integration of electric, calcium, reactive oxygen species and hydraulic signals during rapid systemic signaling in plants. Plant J. 2021:107(1):7–20.

Fichman Y, Zandalinas SI, Peck S, Luan S, and Mittler R. HPCA1 is required for systemic reactive oxygen species and calcium cell-to-cell signaling and plant acclimation to stress. Plant Cell. 2022:34(11):4453–4471.

Figueiredo A, Monteiro F, and Sebastiana M. Subtilisin-like proteases in plantâ€“pathogen recognition and immune priming: a perspective. Front Plant Sci. 2014:5:739.

Fischer, G., Nachtergaele, O. F, Velthuizen V, T. H, Chiozza, F., Franceschini, G., et al. Global Agro-Ecological Zones v4 – Model documentation (Food & Agriculture Org.).

Hakla HR, Sharma S, Urfan M, Mandlik R, Kumawat S, Rajput P, Khajuria B, Chowdhary R, Deshmukh R, Roychowdhury R, et al. Genome-Wide Association Study (GWAS) for Identifying SNPs and Genes Related to Phosphate-Induced Phenotypic Traits in Tomato (Solanum lycopersicum L.). Plants. 2024:13(3). 10.3390/plants13030457

Hasanuzzaman M, Bhuyan MHMB, Anee TI, Parvin K, Nahar K, Mahmud JA, and Fujita M. Regulation of Ascorbate-Glutathione Pathway in Mitigating Oxidative Damage in Plants under Abiotic Stress. Antioxidants (Basel). 2019:8(9). 10.3390/antiox8090384

Hassani A, Azapagic A, and Shokri N. Global predictions of primary soil salinization under changing climate in the 21st century. Nat Commun. 2021:12(1):6663.

Jiang C, Belfield EJ, Cao Y, Smith JAC, and Harberd NP. An Arabidopsis soil-salinity-tolerance mutation confers ethylene-mediated enhancement of sodium/potassium homeostasis. Plant Cell. 2013:25(9):3535–3552.

Julkowska MM, Koevoets IT, Mol S, Hoefsloot H, Feron R, Tester MA, Keurentjes JJB, Korte A, Haring MA, de Boer G-J, et al. Genetic Components of Root Architecture Remodeling in Response to Salt Stress. Plant Cell. 2017:29(12):3198–3213.

Kang HM, Sul JH, Service SK, Zaitlen NA, Kong S-Y, Freimer NB, Sabatti C, and Eskin E. Variance component model to account for sample structure in genome-wide association studies. Nat Genet. 2010:42(4):348–354.

Kassambara A. ggpubr:’ggplot2’Based Publication Ready Plots. R package version 0.4. 0.] https://cran-r-projectorg/web/packages/ggpubr/indes.

Keitt T. Builds Color Tables [R package colorRamps version 2.3.4]. https://cran.r-project.org/web/packages/colorRamps/index.html. Retrieved June 5, 2024

Kim D, Paggi JM, Park C, Bennett C, and Salzberg SL. Graph-based genome alignment and genotyping with HISAT2 and HISAT-genotype. Nat Biotechnol. 2019:37(8):907–915.

Kovach MJ and McCouch SR. Leveraging natural diversity: back through the bottleneck. Curr Opin Plant Biol. 2008:11(2):193–200.

Langfelder P and Horvath S. WGCNA: an R package for weighted correlation network analysis. BMC Bioinformatics. 2008:9:559.

Lee MH and Hwang I. Adaptor proteins in protein trafficking between endomembrane compartments in plants. J Plant Biol. 2014:57(5):265–273.

Liao Y, Smyth GK, and Shi W. featureCounts: an efficient general purpose program for assigning sequence reads to genomic features. Bioinformatics. 2014:30(7):923–930.

Li H. Aligning sequence reads, clone sequences and assembly contigs with BWA-MEM. arXiv [q-bioGN]. 2013.

Li N, He Q, Wang J, Wang B, Zhao J, Huang S, Yang T, Tang Y, Yang S, Aisimutuola P, et al. Super-pangenome analyses highlight genomic diversity and structural variation across wild and cultivated tomato species. Nat Genet. 2023:55(5):852–860.

Lobet G, Pagès L, and Draye X. A novel image-analysis toolbox enabling quantitative analysis of root system architecture. Plant Physiol. 2011:157(1):29–39.

Love M, Anders S, and Huber W. Differential analysis of count data – the DESeq2 package. 2013.

Manaa A, Ben Ahmed H, Valot B, Bouchet J-P, Aschi-Smiti S, Causse M, and Faurobert M. Salt and genotype impact on plant physiology and root proteome variations in tomato. J Exp Bot. 2011:62(8):2797–2813.

Martínez-Cuenca M-R, Pereira-Dias L, Soler S, López-Serrano L, Alonso D, Calatayud Á, and Díez MJ. Adaptation to Water and Salt Stresses of Solanum pimpinellifolium and Solanum lycopersicum var. cerasiforme. Agronomy (Basel). 2020:10(8):1169.

McKenna A, Hanna M, Banks E, Sivachenko A, Cibulskis K, Kernytsky A, Garimella K, Altshuler D, Gabriel S, Daly M, et al. The Genome Analysis Toolkit: a MapReduce framework for analyzing next-generation DNA sequencing data. Genome Res. 2010:20(9):1297–1303.

Melino V and Tester M. Salt-Tolerant Crops: Time to Deliver. Annu Rev Plant Biol. 2023:74:671–696.

Morton M, Fiene G, Ahmed HI, Rey E, Abrouk M, Angel Y, Johansen K, Saber NO, Malbeteau Y, Al-Mashharawi S, et al. Deciphering Salt Stress Responses in Solanum pimpinellifolium through High-Throughput Phenotyping. bioRxiv. 2023:2023.08.15.553433. 10.1101/2023.08.15.553433

Munns R and Tester M. Mechanisms of salinity tolerance. Annu Rev Plant Biol. 2008:59:651–681.

Myers RJ Jr, Zandalinas SI, and Mittler R. Live whole-plant detection of rapidly accumulating reactive Oxygen Species following applied stress in Arabidopsis thaliana. Methods Mol Biol. 2023:2642:387–401.

Neuteboom LW, Veth-Tello LM, Clijdesdale OR, Hooykaas PJ, and van der Zaal BJ. A novel subtilisin-like protease gene from Arabidopsis thaliana is expressed at sites of lateral root emergence. DNA Res. 1999:6(1):13–19.

Ohlsson JA, Leong JX, Elander PH, Dauphinee AN, Ballhaus F, Johansson J, Lommel M, Hofmann G, Betnér S, Sandgren M, et al. SPIRO – the automated Petri plate imaging platform designed by biologists, for biologists. bioRxiv. 2021:2021.03.15.435343. 10.1101/2021.03.15.435343

Pailles Y, Awlia M, Julkowska M, Passone L, Zemmouri K, Negrão S, Schmöckel SM, and Tester M. Diverse Traits Contribute to Salinity Tolerance of Wild Tomato Seedlings from the Galapagos Islands. Plant Physiol. 2020:182(1):534–546.

Paran I and van der Knaap E. Genetic and molecular regulation of fruit and plant domestication traits in tomato and pepper. J Exp Bot. 2007:58(14):3841–3852.

Pease JB, Haak DC, Hahn MW, and Moyle LC. Phylogenomics Reveals Three Sources of Adaptive Variation during a Rapid Radiation. PLoS Biol. 2016:14(2):e1002379.

Peng J, Li Z, Wen X, Li W, Shi H, Yang L, Zhu H, and Guo H. Salt-induced stabilization of EIN3/EIL1 confers salinity tolerance by deterring ROS accumulation in Arabidopsis. PLoS Genet. 2014:10(10):e1004664.

Rao ES, Kadirvel P, Symonds RC, Geethanjali S, Thontadarya RN, and Ebert AW. Variations in DREB1A and VP1.1 Genes Show Association with Salt Tolerance Traits in Wild Tomato (Solanum pimpinellifolium). PLoS One. 2015:10(7):e0132535.

Razali R, Bougouffa S, Morton MJL, Lightfoot DJ, Alam I, Essack M, Arold ST, Kamau AA, Schmöckel SM, Pailles Y, et al. The genome sequence of the wild tomato Solanum pimpinellifolium provides insights into salinity tolerance. Front Plant Sci. 2018:9:1402.

Richards LA. Diagnosis and Improvement of Saline and Alkali Soils (United States Department of Agriculture).

Rodriguez M, Scintu A, Posadinu CM, Xu Y, Nguyen CV, Sun H, Bitocchi E, Bellucci E, Papa R, Fei Z, et al. GWAS Based on RNA-Seq SNPs and High-Throughput Phenotyping Combined with Climatic Data Highlights the Reservoir of Valuable Genetic Diversity in Regional Tomato Landraces. Genes. 2020:11(11). 10.3390/genes11111387

Roșca M, Mihalache G, and Stoleru V. Tomato responses to salinity stress: From morphological traits to genetic changes. Front Plant Sci. 2023:14:1118383.

Růzicka K, Ljung K, Vanneste S, Podhorská R, Beeckman T, Friml J, and Benková E. Ethylene regulates root growth through effects on auxin biosynthesis and transport-dependent auxin distribution. Plant Cell. 2007:19(7):2197–2212.

Sarfatti M, Abu-Abied M, Katan J, and Zamir D. RFLP mapping of I1, a new locus in tomato conferring resistance against Fusarium oxysporum f. sp. lycopersici race 1. Theor Appl Genet. 1991:82(1):22–26.

Shalata A and Neumann PM. Exogenous ascorbic acid (vitamin C) increases resistance to salt stress and reduces lipid peroxidation. J Exp Bot. 2001:52(364):2207–2211.

Shevelev IV and Hübscher U. The 3’ 5’ exonucleases. Nat Rev Mol Cell Biol. 2002:3(5):364–376.

Sievert C. Interactive Web-Based Data Visualization with R, plotly, and shiny (CRC Press).

Sousa B, Rodrigues F, Soares C, Martins M, Azenha M, Lino-Neto T, Santos C, Cunha A, and Fidalgo F. Impact of Combined Heat and Salt Stresses on Tomato Plants-Insights into Nutrient Uptake and Redox Homeostasis. Antioxidants (Basel). 2022:11(3). 10.3390/antiox11030478

Steppuhn H and Raney JP. Emergence, height, and yield of canola and barley grown in saline root zones. Can J Plant Sci. 2005:85(4):815–827.

Street IH, Aman S, Zubo Y, Ramzan A, Wang X, Shakeel SN, Kieber JJ, and Schaller GE. Ethylene inhibits cell proliferation of the Arabidopsis root meristem. Plant Physiol. 2015:169(1):338–350.

Takagi H, Abe A, Yoshida K, Kosugi S, Natsume S, Mitsuoka C, Uemura A, Utsushi H, Tamiru M, Takuno S, et al. QTL-seq: rapid mapping of quantitative trait loci in rice by whole genome resequencing of DNA from two bulked populations. Plant J. 2013:74(1):174–183.

Thabet SG, Moursi YS, Sallam A, Karam MA, and Alqudah AM. Genetic associations uncover candidate SNP markers and genes associated with salt tolerance during seedling developmental phase in barley. Environ Exp Bot. 2021:188:104499.

Thole JM, Beisner ER, Liu J, Venkova SV, and Strader LC. Abscisic acid regulates root elongation through the activities of auxin and ethylene in Arabidopsis thaliana. G3 (Bethesda). 2014:4(7):1259–1274.

Tripodi P, Soler S, Campanelli G, Díez MJ, Esposito S, Sestili S, Figàs MR, Leteo F, Casanova C, Platani C, et al. Genome wide association mapping for agronomic, fruit quality, and root architectural traits in tomato under organic farming conditions. BMC Plant Biol. 2021:21(1):481.

Voiniciuc C. Modern mannan: a hemicellulose’s journey. New Phytol. 2022:234(4):1175–1184.

Wang J, Yu Y, Zhang Z, Quan R, Zhang H, Ma L, Deng XW, and Huang R. Arabidopsis CSN5B interacts with VTC1 and modulates ascorbic acid synthesis. Plant Cell. 2013:25(2):625–636.

Wang J and Zhang Z. GAPIT Version 3: Boosting Power and Accuracy for Genomic Association and Prediction. Genomics Proteomics Bioinformatics. 2021:19(4):629–640.

Wang X, Gao L, Jiao C, Stravoravdis S, Hosmani PS, Saha S, Zhang J, Mainiero S, Strickler SR, Catala C, et al. Genome of Solanum pimpinellifolium provides insights into structural variants during tomato breeding. Nat Commun. 2020a:11(1):5817.

Wang Y, Ji Y, Fu Y, and Guo H. Ethylene-induced microtubule reorientation is essential for fast inhibition of root elongation in Arabidopsis. J Integr Plant Biol. 2018:60(9):864–877.

Wang Z, Hong Y, Zhu G, Li Y, Niu Q, Yao J, Hua K, Bai J, Zhu Y, Shi H, et al. Loss of salt tolerance during tomato domestication conferred by variation in a Na+ /K+ transporter. EMBO J. 2020b:39(10):e103256.

Website. 10.1104/pp.111.192088

Wei T and Simko V. R package “corrplot”: Visualization of a Correlation Matrix (Version 0.84). Retrived from, https://githubcom/taiyun/corrplot. 2017.

Wickham H. ggplot2: Elegant Graphics for Data Analysis (Springer).

Wickham H. reshape2: flexibly reshape data: a reboot of the reshape package. R package version. 2020.

Wickham H, Averick M, Bryan J, Chang W, McGowan L, François R, Grolemund G, Hayes A, Henry L, Hester J, et al. Welcome to the tidyverse. J Open Source Softw. 2019:4(43):1686.

Wilke CO. cowplot: streamlined plot theme and plot annotations for “ggplot2.” CRAN Repos. 2016.

Yang H, Du T, Mao X, Ding R, and Shukla MK. A comprehensive method of evaluating the impact of drought and salt stress on tomato growth and fruit quality based on EPIC growth model. Agric Water Manage. 2019:213:116–127.

Yassin TE. Inheritance of resistance to leaf curl virus disease in a cross between tomato (Lycopersicon esculentum Mill.) and currant tomato (L. pimpinellifolium (Jusl.) Mill.). J Agric Sci. 1985:105(3):659–661.

Ye J, Wang X, Wang W, Yu H, Ai G, Li C, Sun P, Wang X, Li H, Ouyang B, et al. Genome-wide association study reveals the genetic architecture of 27 agronomic traits in tomato. Plant Physiol. 2021:186(4):2078–2092.

Zhang R, Li B, Zhao Y, Zhu Y, and Li L. An essential role for mannan degradation in both cell growth and secondary cell wall formation. J Exp Bot. 2024:75(5):1407–1420.

Zhang S, Yu H, Wang K, Zheng Z, Liu L, Xu M, Jiao Z, Li R, Liu X, Li J, et al. Detection of major loci associated with the variation of 18 important agronomic traits between Solanum pimpinellifolium and cultivated tomatoes. Plant J. 2018:95(2):312–323.

Zhang Z, Wang J, Zhang R, and Huang R. The ethylene response factor AtERF98 enhances tolerance to salt through the transcriptional activation of ascorbic acid synthesis in Arabidopsis. Plant J. 2012:71(2):273–287.

