## Supplemental Figures for "Ethylene and ROS Signaling Are Key Regulators of Lateral Root Development under Salt Stress in Tomato"

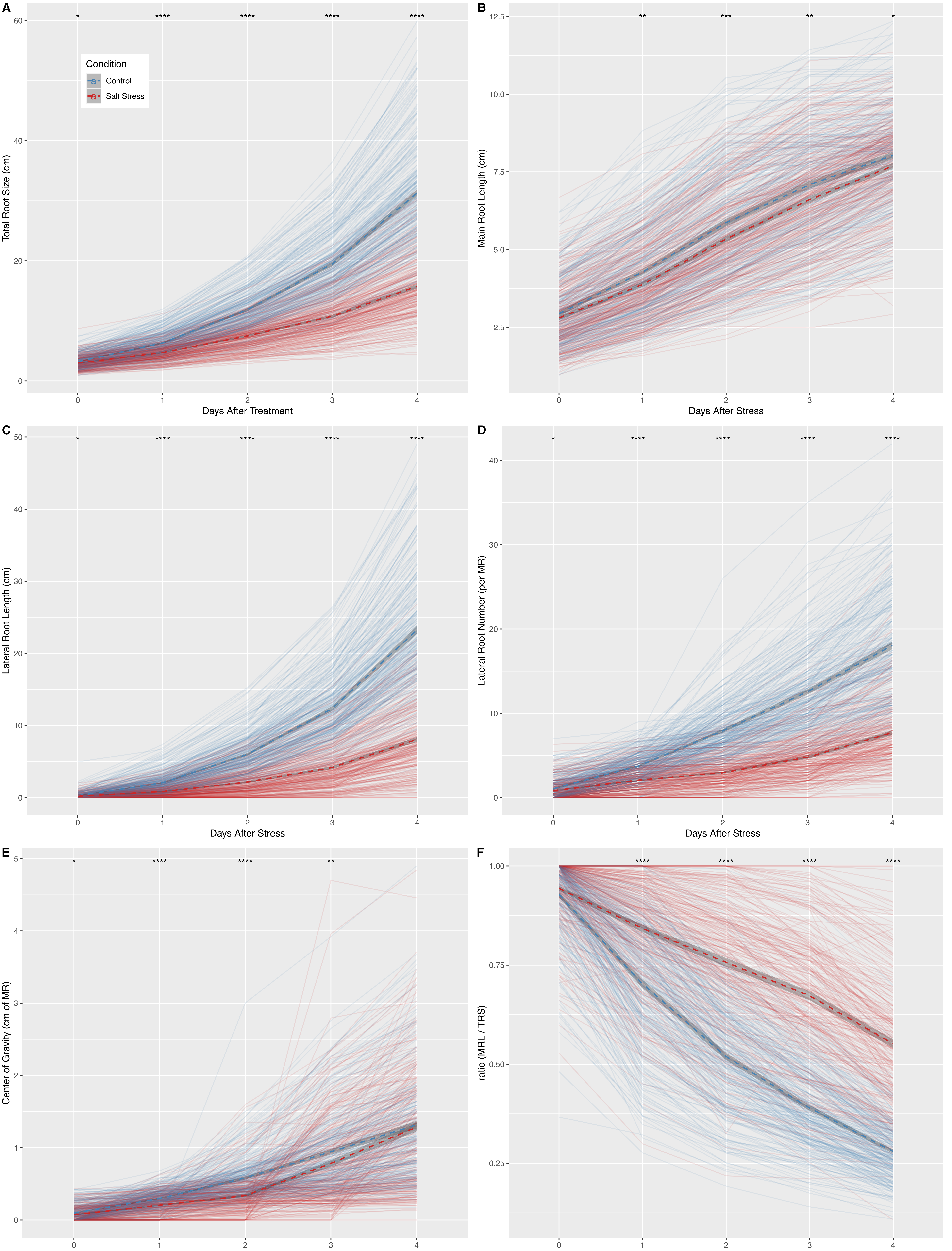


**Figure S1. The natural variation in Root System Architecture components in response to salt stress.** The 220 genotypes of S. pimpinellifolium were germinated on ½ MS agar plates for 4 days and subsequently transferred to media containing 0 or 100 mM NaCl (Control and Salt respectively). The seedlings were imaged every 24 h after transfer for 7 consecutive days. The Root Architecture of seedlings was quantified for the first 4 days after transfer, and inspected for changes throughout time for **(A)** Total Root Size, **(B)** Main Root Length, **(C)** Lateral Root Length, **(D)** Lateral Root Number, **(E)** Center of Gravity (calculated as the position of MR with most Lateral Root Length), and **(F)** ratio of Main Root Length per Total Root Size. The individual lines represent the genotype-specific mean, whereas the dashed lines represent the population average. The *, **, *** and **** represent the significant differences between the treatment at individual time points as tested with two-way ANOVA with p-value < 0.05, 0.01, 0.001 and 0.0001, respectively.

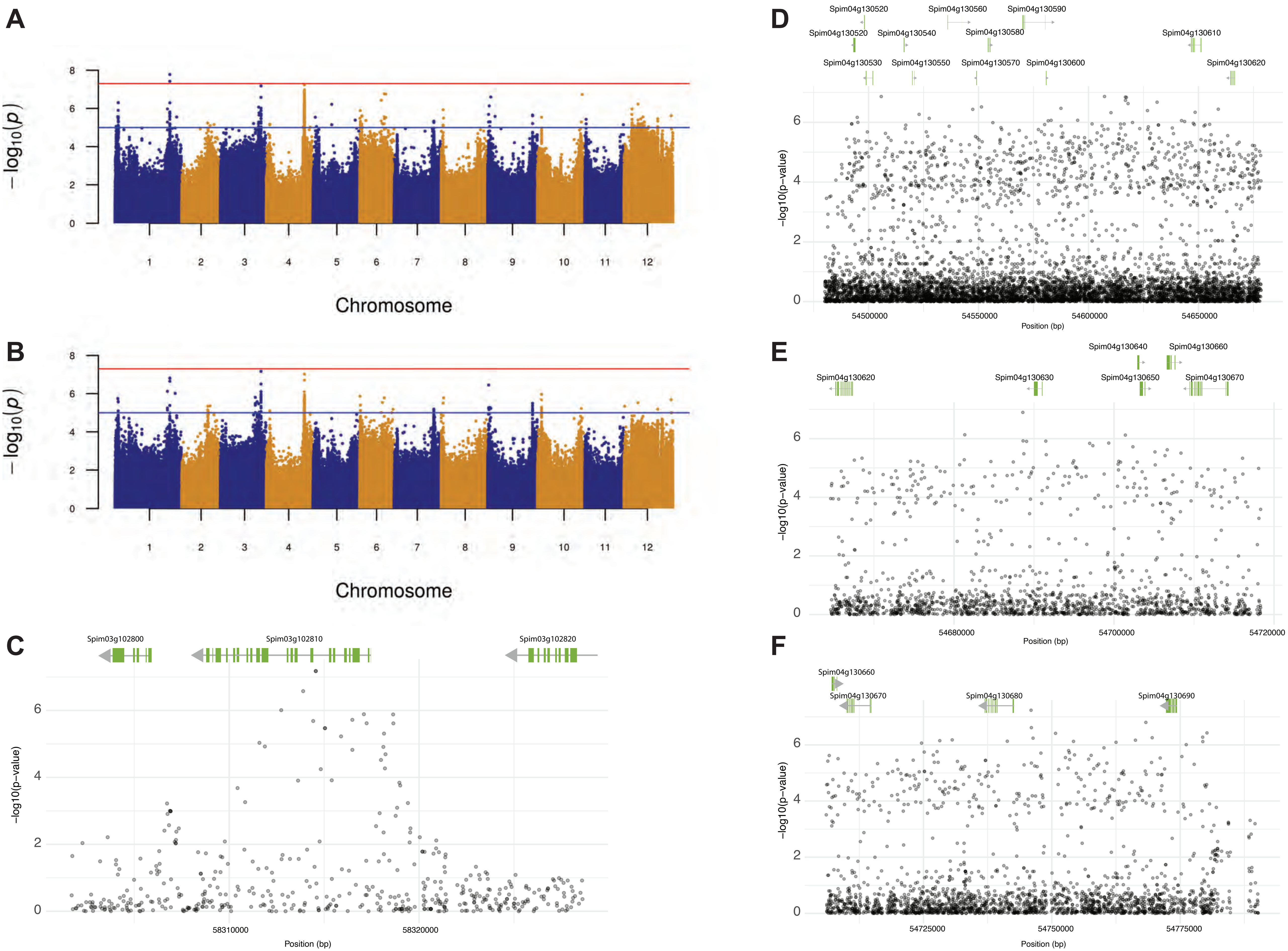


**Figure S2. Salt-induced changes in root biomass distribution between main and lateral roots are associated with loci on Chromosomes 3 and 4.** Genome Wide Association Study was performed on 9M SNPs using standard EMMA-X method. Only SNPs with minor allele frequency > 0.05 were used for plotting the Manhattan plot for associations identified with **(A)** ratio of main root length to total root size at 1 day and **(B)** two days post transfer to salt stress, with pseudo-heritability values of 0.116 and 0.195 respectively. The red line indicates the Bonferroni threshold, whereas the blue line indicates genome-wide suggestive threshold. **(C)** The significantly associated loci were inspected in further detail for the gene coding sequences within the regions with significantly associated SNPs. The region on chromosome 3 contained 29 SNPs with -log10(p-value) > 5. The three individual loci **(D-F)** identified on chromosome 4 contained 240, 39 and 85 SNPs with -log10(p-value) > 5. The association graph below each locus represents the association strength with the ratio of main root length to total root size at 1 day after salt stress imposition.

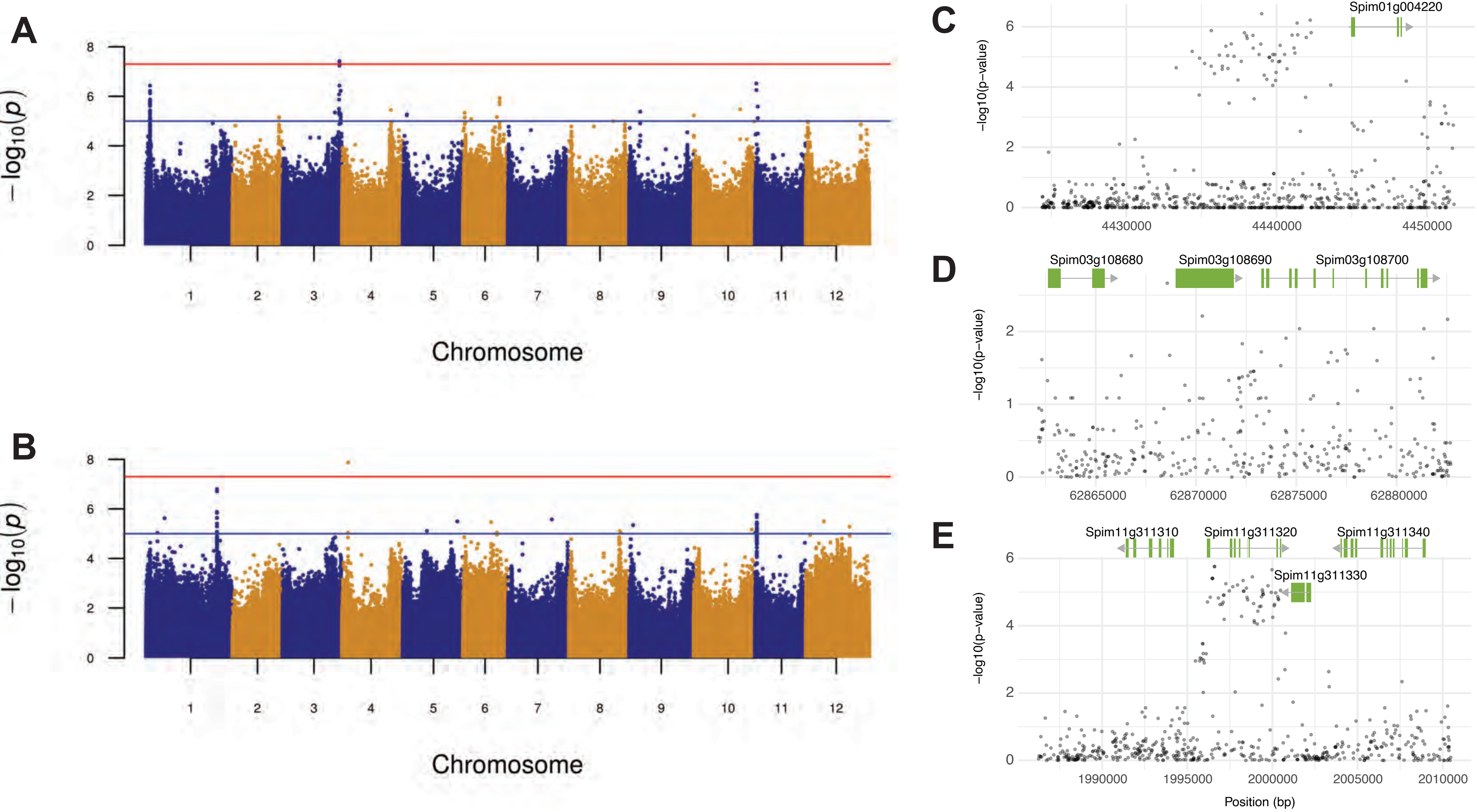


**Figure S3. Salt-induced changes in lateral root development are associated with loci on chromosomes 1, 3 and 10. Genome Wide Association Study was performed on 9M SNPs using standard EMMA-X method.** Only SNPs with minor allele frequency > 0.05 were used for plotting the Manhattan plot for associations identified with **(A)** average lateral root length and **(B)** lateral root number at one day post transfer to salt stress, with pseudo-heritability values of 0.145 and 0.358 respectively. The red line indicates the Bonferroni threshold, whereas the blue line indicates genome-wide suggestive threshold. **(C)** The significantly associated loci were inspected in further detail for the gene coding sequences within the regions with significantly associated SNPs. The region on chromosome 1 contained 27 SNPs, **(D)** the region on chromosome 3 7 SNPs, and **(E)** region on chromosome 11 contained 60 SNPs with -log10(p-value) > 5. The association graph below each locus represents the association strength with the average lateral root length at 1 day after salt stress imposition.

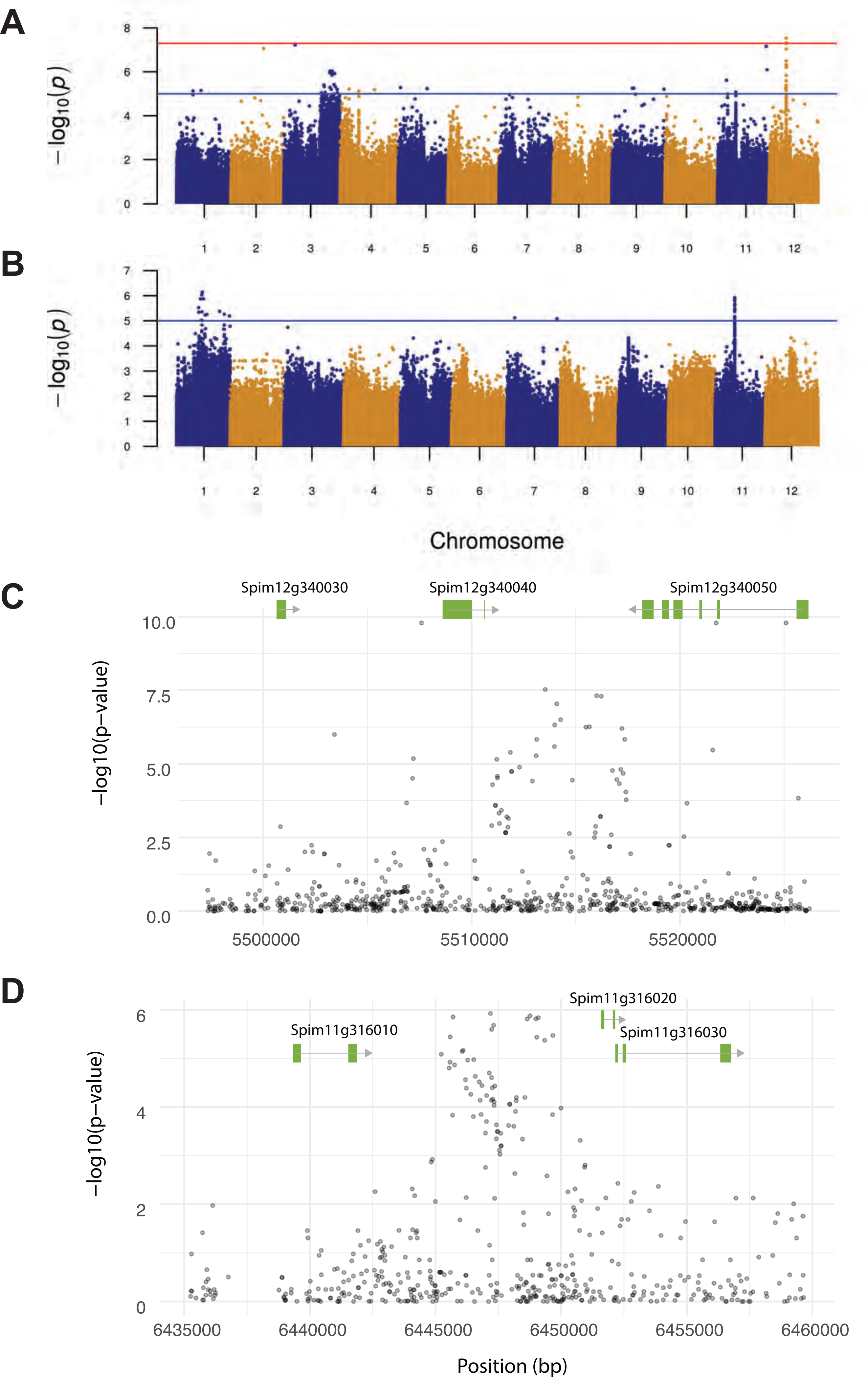


**Figure S4. Emergence of lateral roots and main root growth rate is associated with loci on chromosomes 12 and 22.** Genome Wide Association Study was performed on 9M SNPs using standard EMMA-X method. Only SNPs with minor allele frequency > 0.05 were used for plotting the Manhattan plot for associations identified with **(A)** lateral root emergence and **(B)** main root growth rate under salt stress, with pseudo-heritability values of 0.855 and 0.628 respectively. The red line indicates the Bonferroni threshold, whereas the blue line indicates genome-wide suggestive threshold. **(C)** The significantly associated loci were inspected in further detail for the gene coding sequences within the regions with significantly associated SNPs. The region on chromosome 12 contained 16 SNPs, whereas **(D)** the region on chromosome 11 contained 15 SNPs -log10(p-value) > 5. The association graph below each locus represents the association strength with the trait initially associated with that locus.

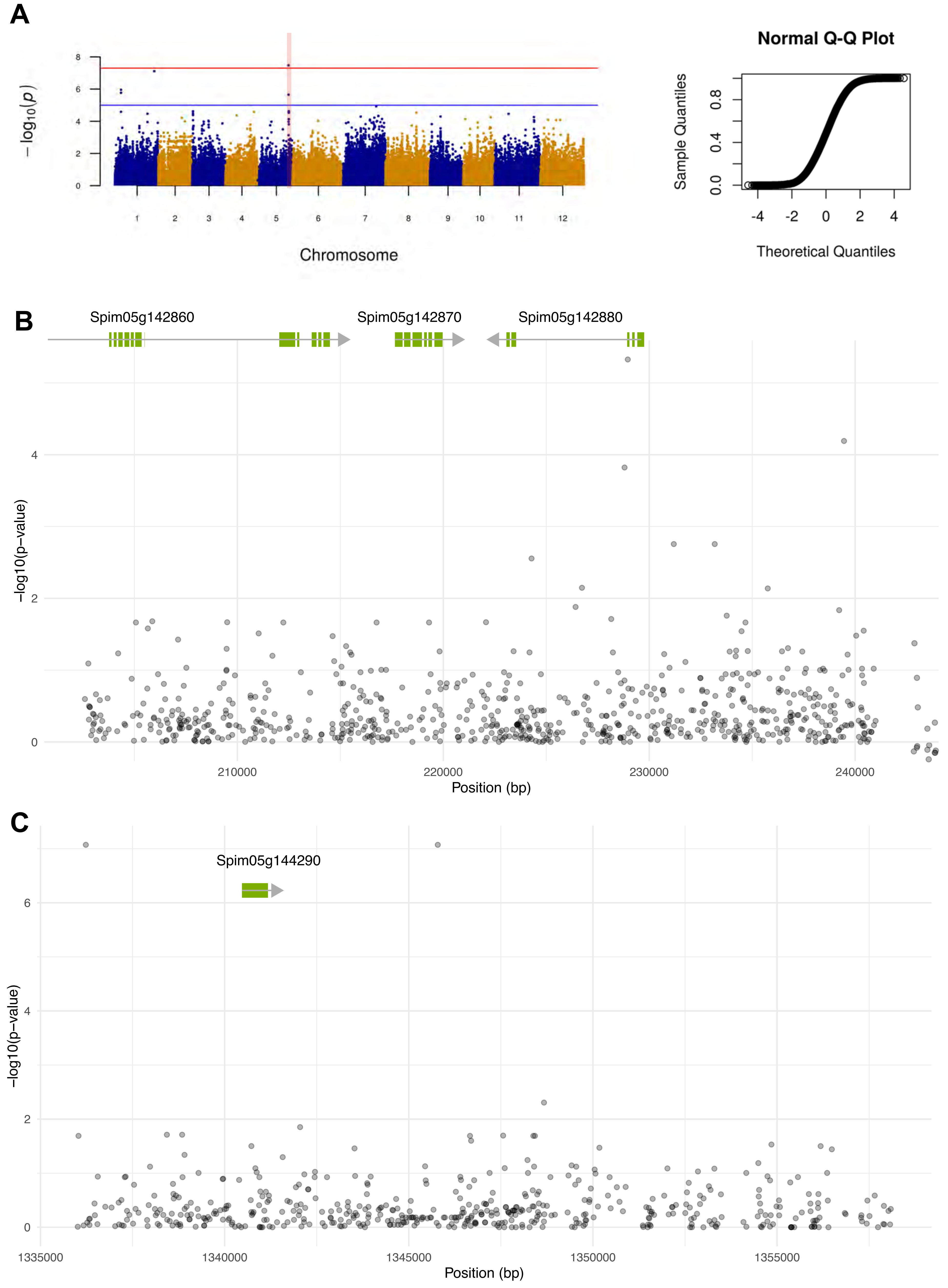


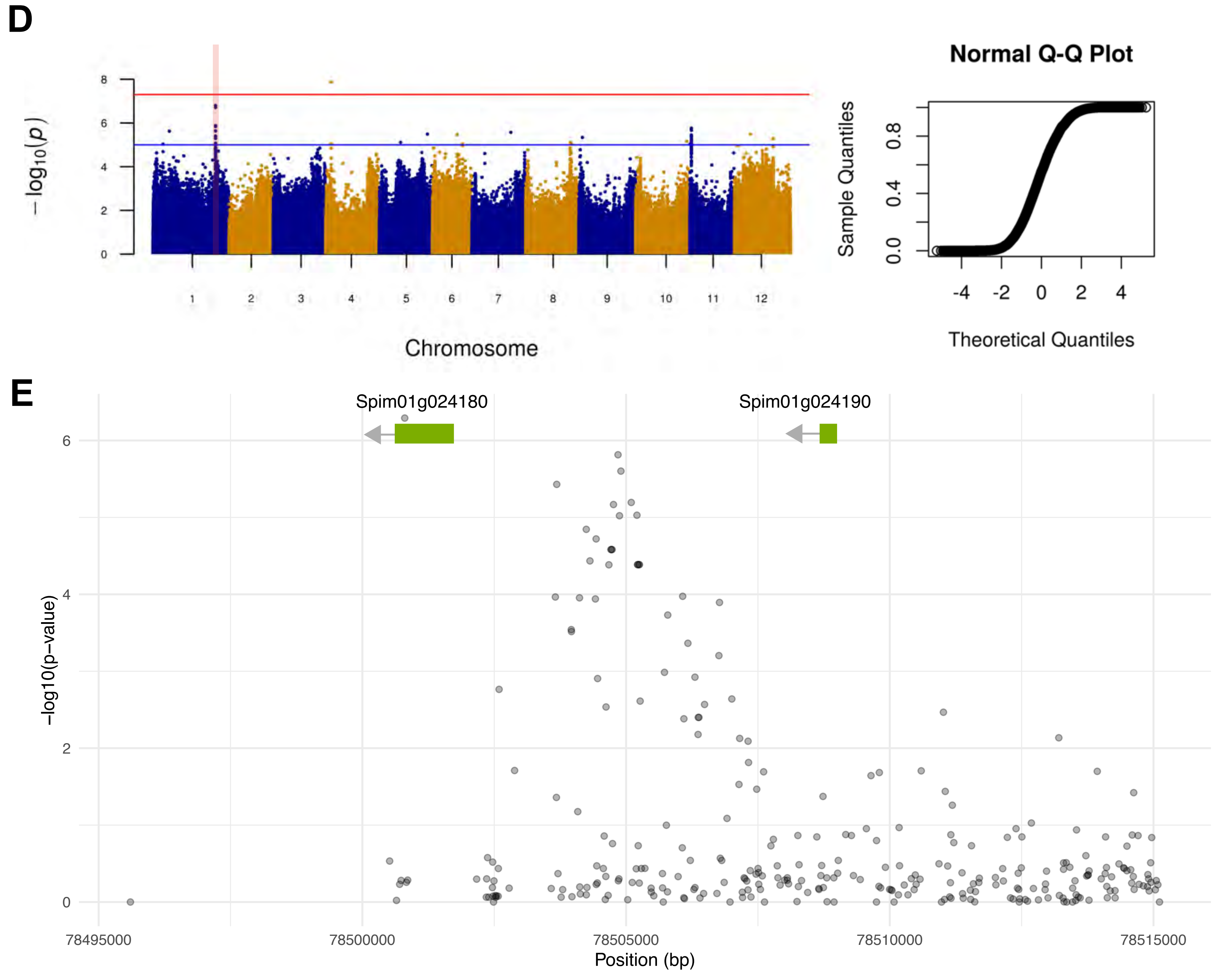


**Figure S5. High-confidence loci identified through GWAS. Genome Wide Association Study was performed on 9M SNPs using the standard EMMA-X method.** Only SNPs with minor allele frequency > 0.05 were used for plotting the Manhattan plot for associations identified with **(A)** lateral root number at 2 days after transfer to control conditions. The significantly associated loci on chromosome 5, consisting of **(B)** 15 and **(C)** 11 SNPs were inspected in further detail. **(D)** Associations with the number of lateral roots were identified on chromosome 1, and **(E)** the locus consisting of 14 highly associated SNPs has been inspected in more detail. The red line on Manhattan plots indicates the Bonferroni threshold, whereas the blue line indicates genome-wide suggestive threshold. The association graph below each locus represents the association strength with the trait initially associated with that locus.

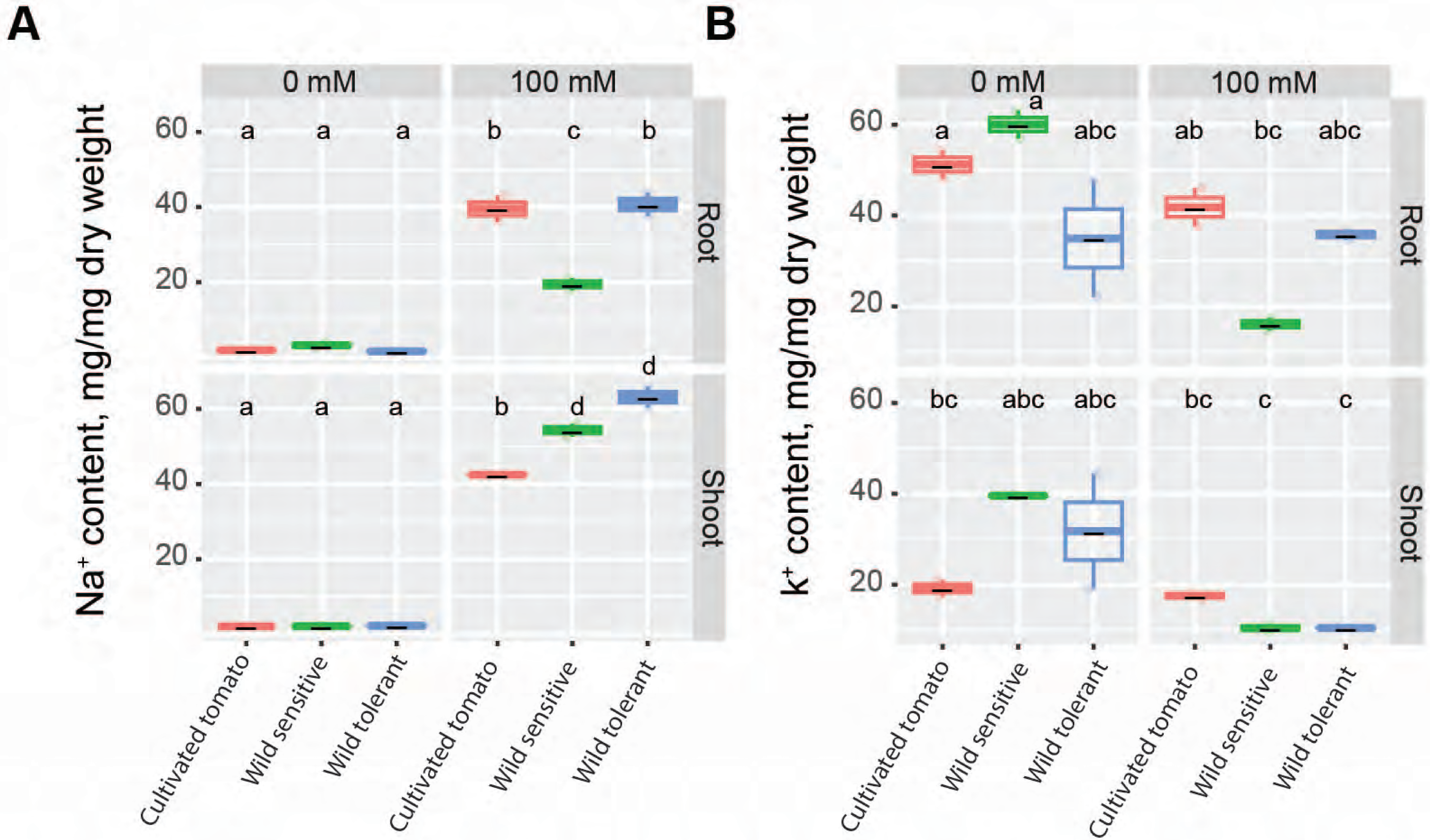


**Figure S6. Wild tomatoes accumulate higher levels of Na+ in the shoot compared to cultivated tomato. (A)** Na+ and **(B)** K+ contents of root and shoot of different accessions after 10 days on treatment plates. Each dot represents an individual replicate per accession. Statistical analysis was done by comparison of the means for all pairs using Tukey HSD test. Levels not connected by the same letter are significantly different ($P < 0.05$).

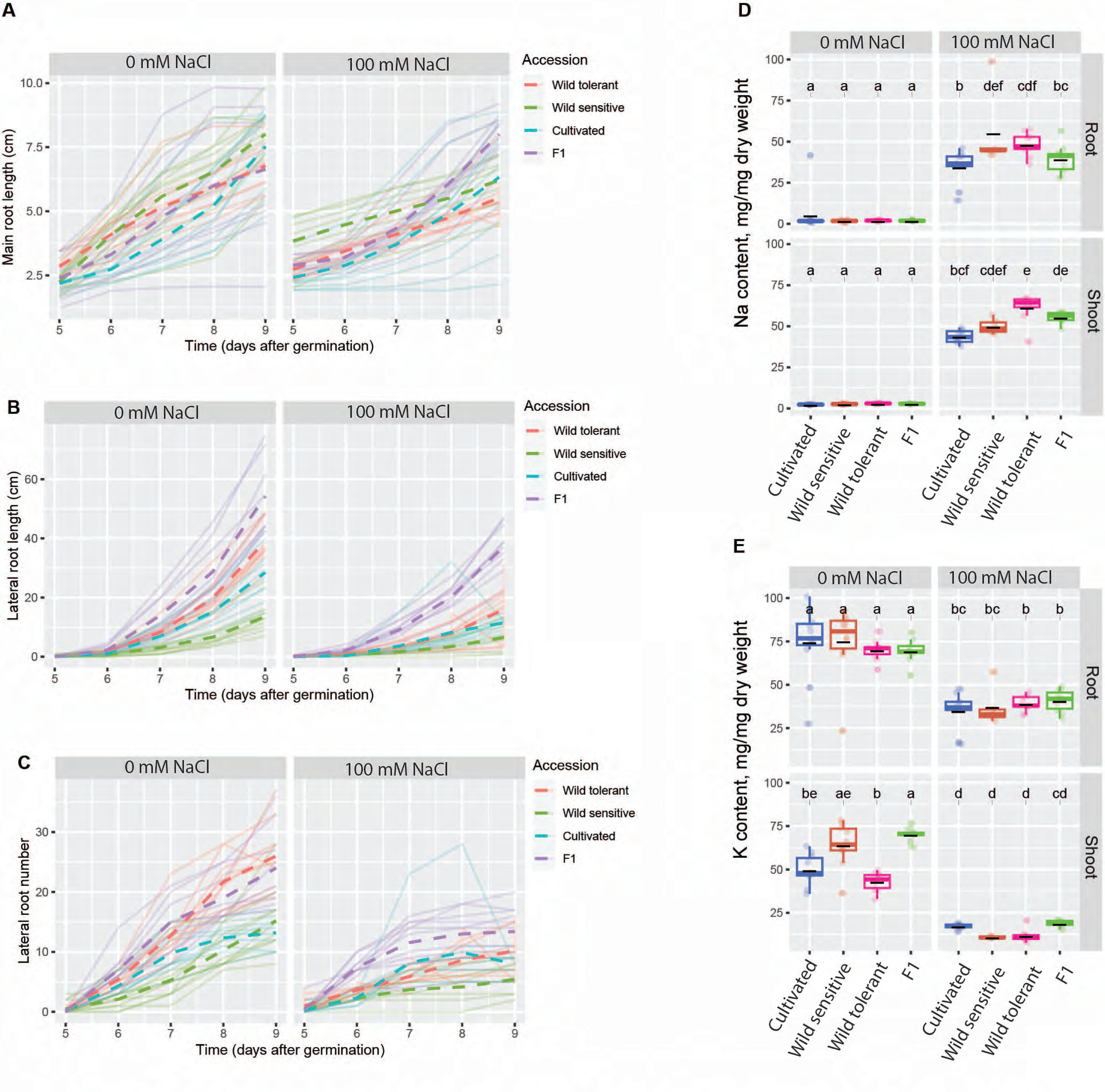


**Figure S7. F1 individuals accumulate less Na+. (A-C)** Root system architecture of F1 individuals, their corresponding parents, i.e., cultivated and wild tolerant tomatoes along with wild sensitive tomatoes were investigated with and without 100 mM NaCl. **(D-E)** Na+ and K+ contents of root and shoot of different accessions after 10 days on treatment plates. Each dot represents individual replicas per accession. Statistical analysis was done by comparison of the means for all pairs using Tukey HSD test in all graphs. Levels not connected by the same letter are significantly different ($P < 0.05$).

**Figure S8. Remaining bulk comparisons.** Displayed here are the additional BSA results that were not shown in Figure 4.

Low LRL / low LR.no vs. Low LRL / high LR.no

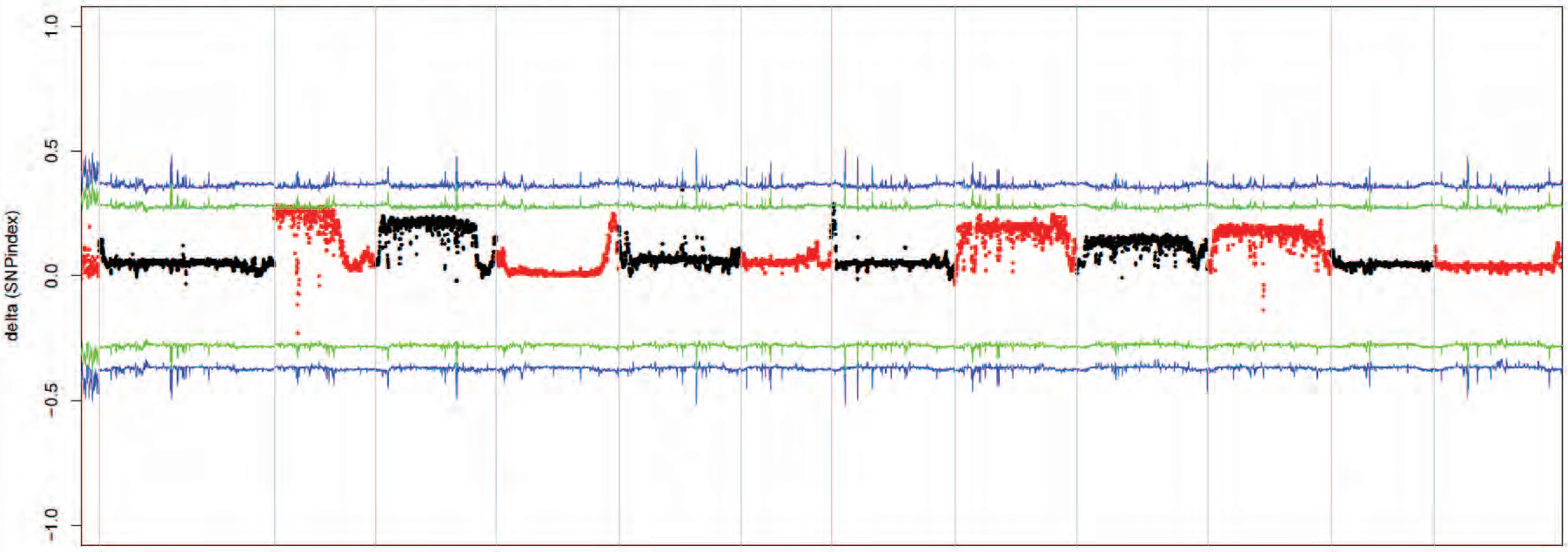


Low LRL / low LR.no vs. High LRL / low LR.no

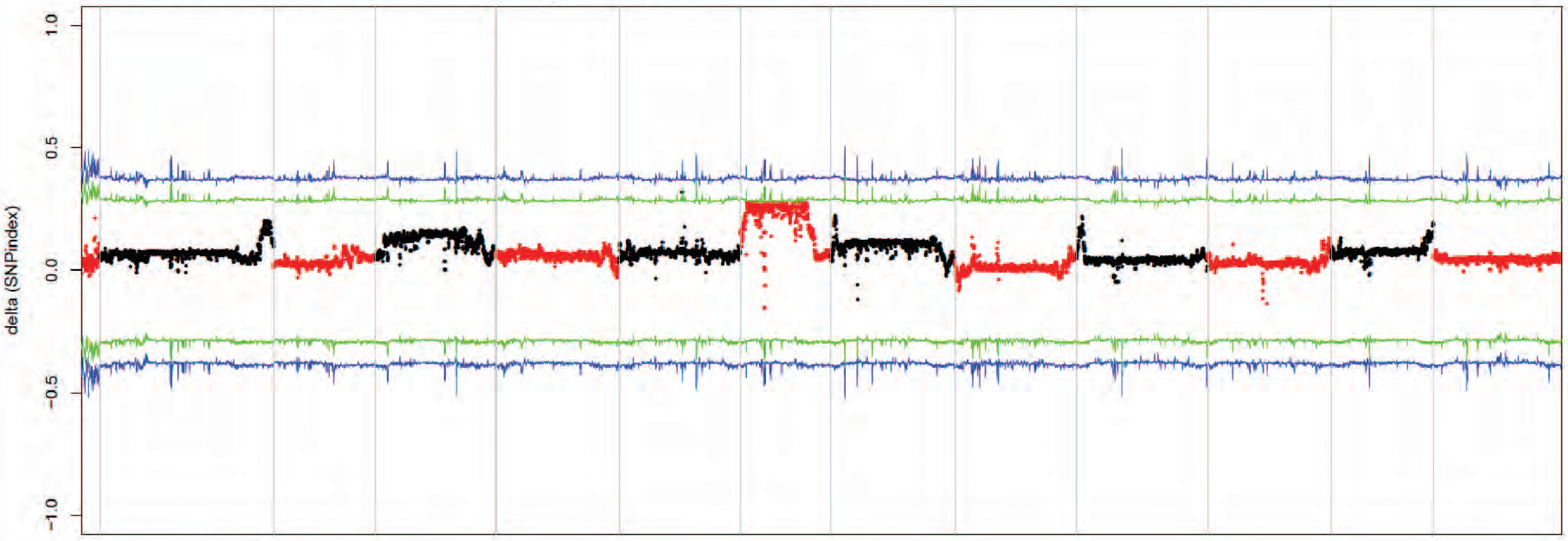


Lateral root lenght;
Low LRL / low LR.no+ Low LRL / high LR.no vs. High LRL / low LR.no + High LRL / High LR.no

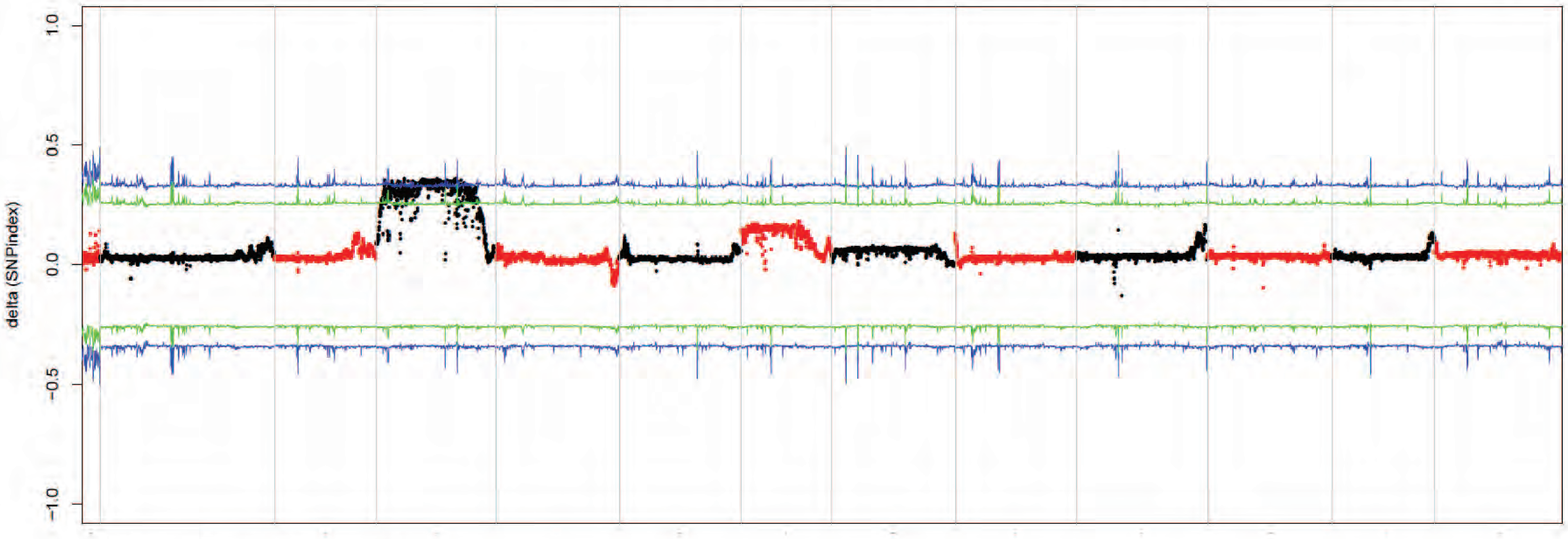


Lateral root number;
Low LRL / low LR.no + High LRL / low LR.no vs. Low LRL / high LR.no + High LRL / High LR.no

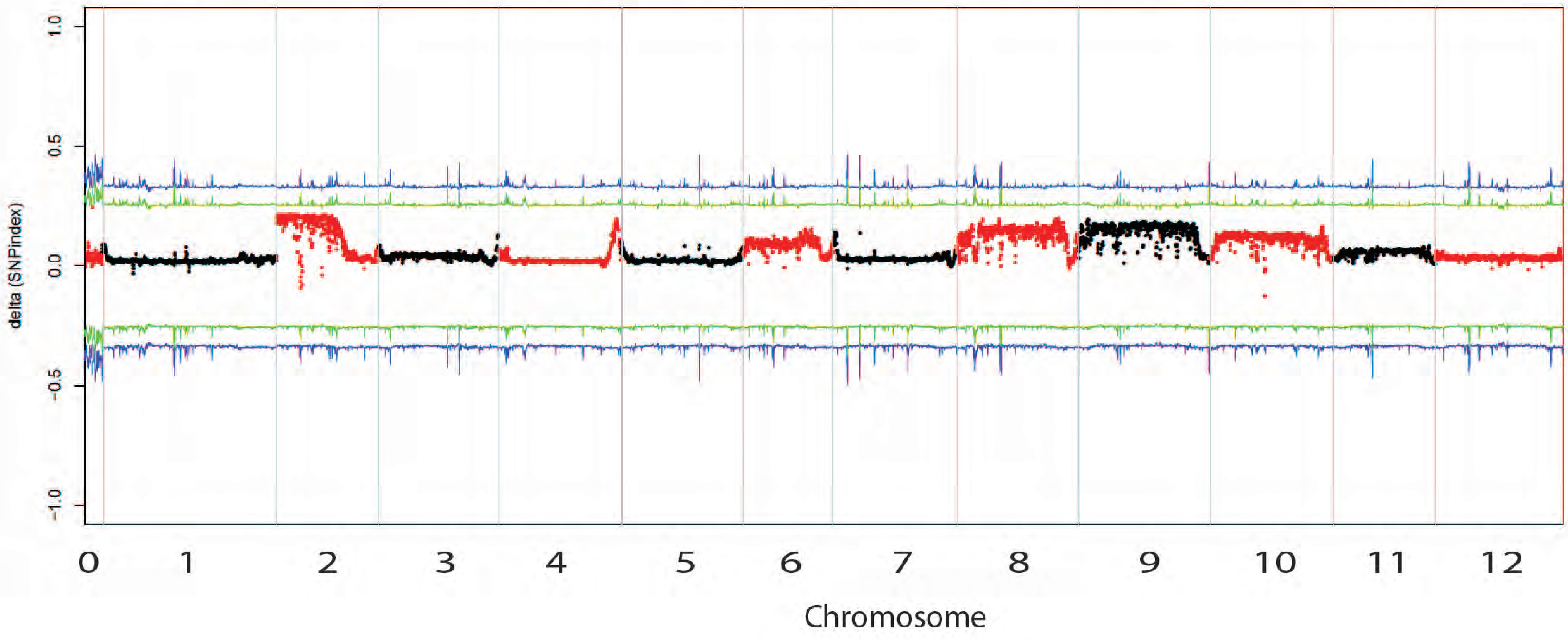

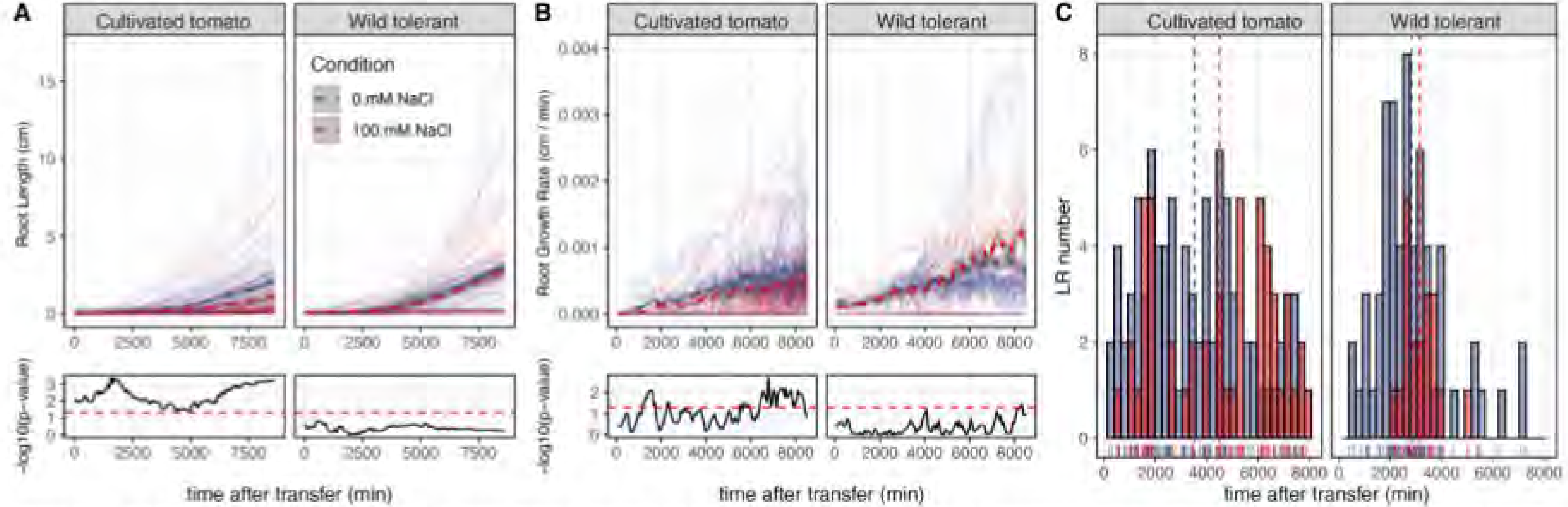


**Figure 8. Wild tolerant and cultivated tomatoes show variation in lateral root elongation and emergence under the salt stress.** (A) Lateral root length, (B) lateral root length growth rate, and (C) lateral root number are shown for the two tomato accessions grown in agar plate under 100 mM NaCl. The plates were imaged using SPIRO setup in a 30-min interval for one week. Individual transparent lines indicate the growth trajectory of individual lateral root, dashed line indicates the growth average per genotype per condition, whereas the shaded area indicates the standard deviation. The differences between Control and Salt conditions were tested within each genotype using t-test and the -log10(p-value) graphs are displayed under each graph. The red dashed line represents a threshold corresponding to p-value of 0.05. The line above the threshold indicates significant difference between Control and Salt Stress conditions. The growth dynamics of lateral roots were measured over two experimental replicates for 4 biological replicates per genotype per condition.

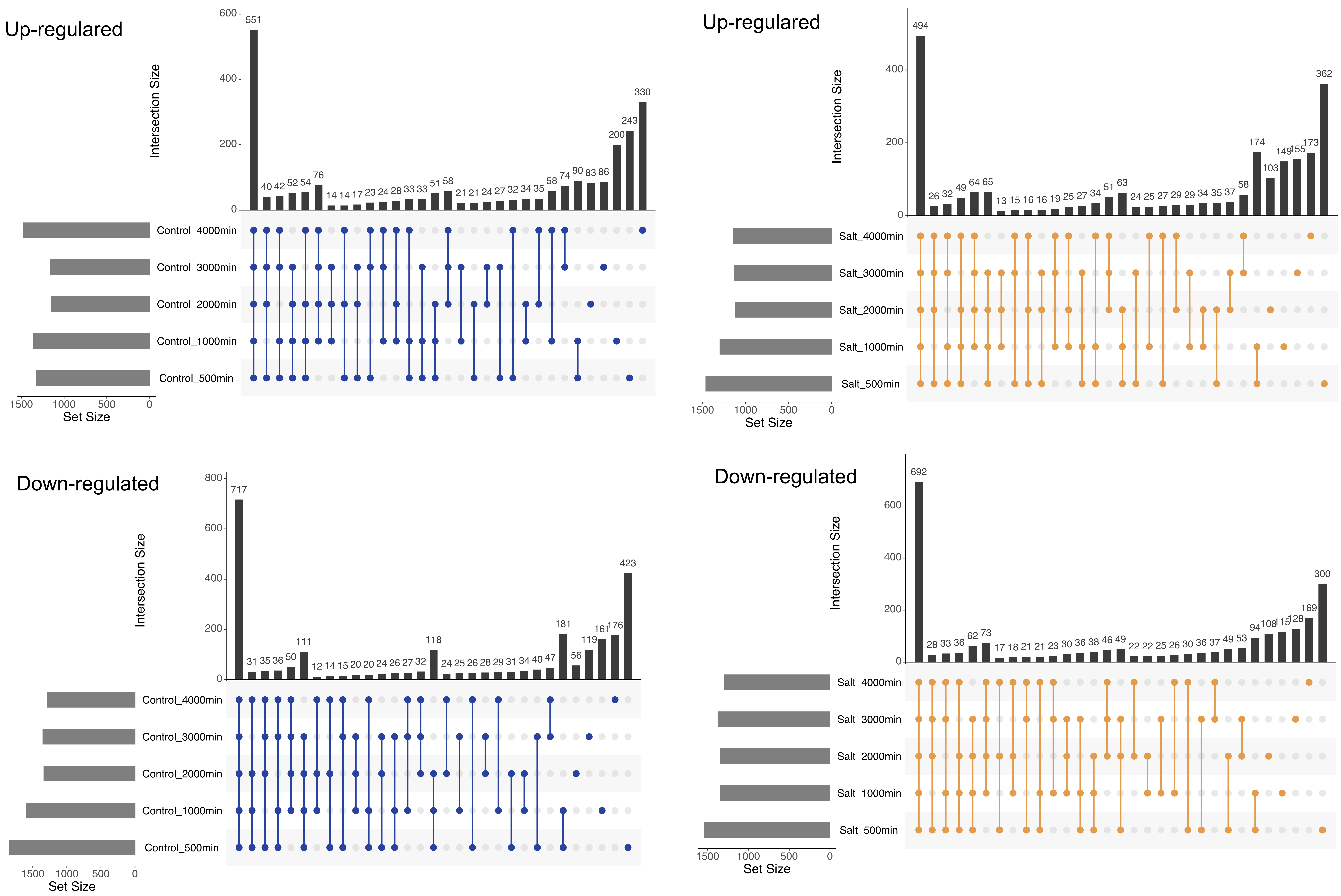


**Figure S10. Intersection of DEGs between WT and CT under Salt and Control conditions.** The upset plots highlight the intersection of numbers of DEGs between wild tolerant and cultivated tomato samples across five timepoints under salt and control conditions, respectively. The panel with dark blue dots represents samples under the control condition, while the panel with brown dots are Salt samples. The direction of arrows within each panel indicates whether the depicted genes are up- or down-regulated.

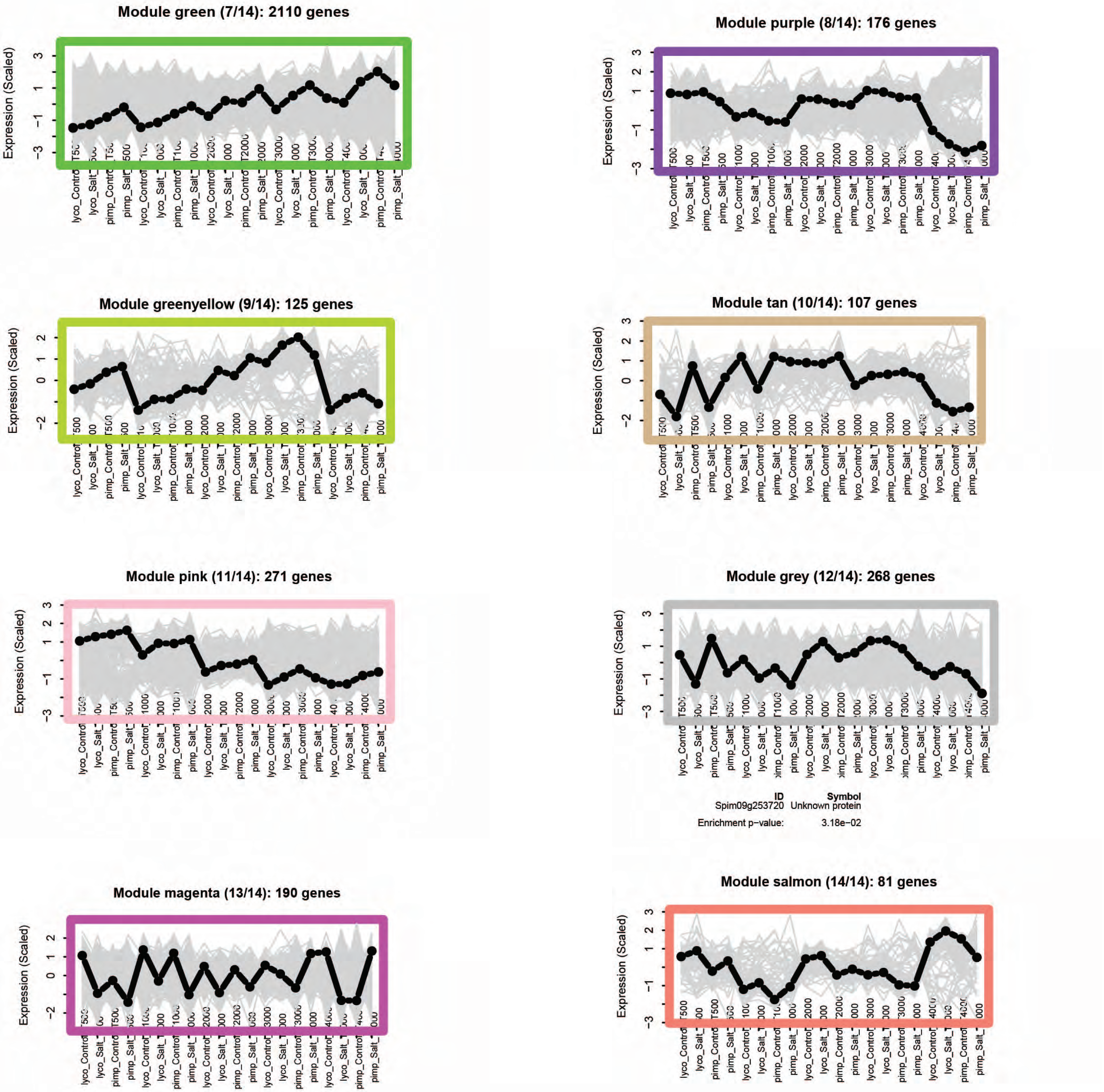


**Figure S11. Co-expression network analysis.** The rest of WGCNA co-expression modules are shown here, module 7 to 14.

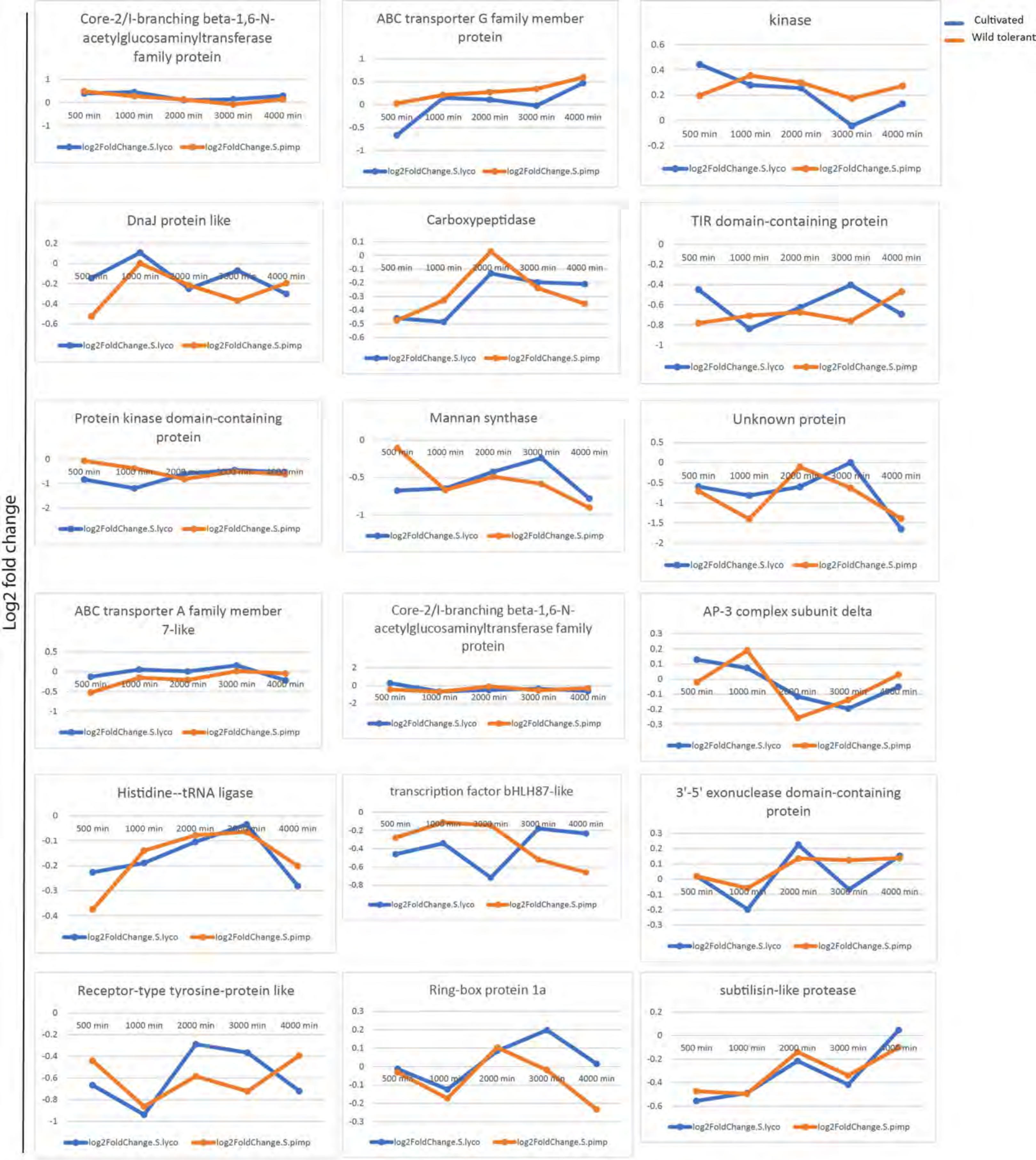


**Figure S12. Gene expression profiles over time for genes identified by overlapping GWAS and BSA analyses in Table 1.** Log2 fold changes are shown for the 18 genes identified by cross-referencing the GWAS and BSA, across all developmental time points for the two accessions.

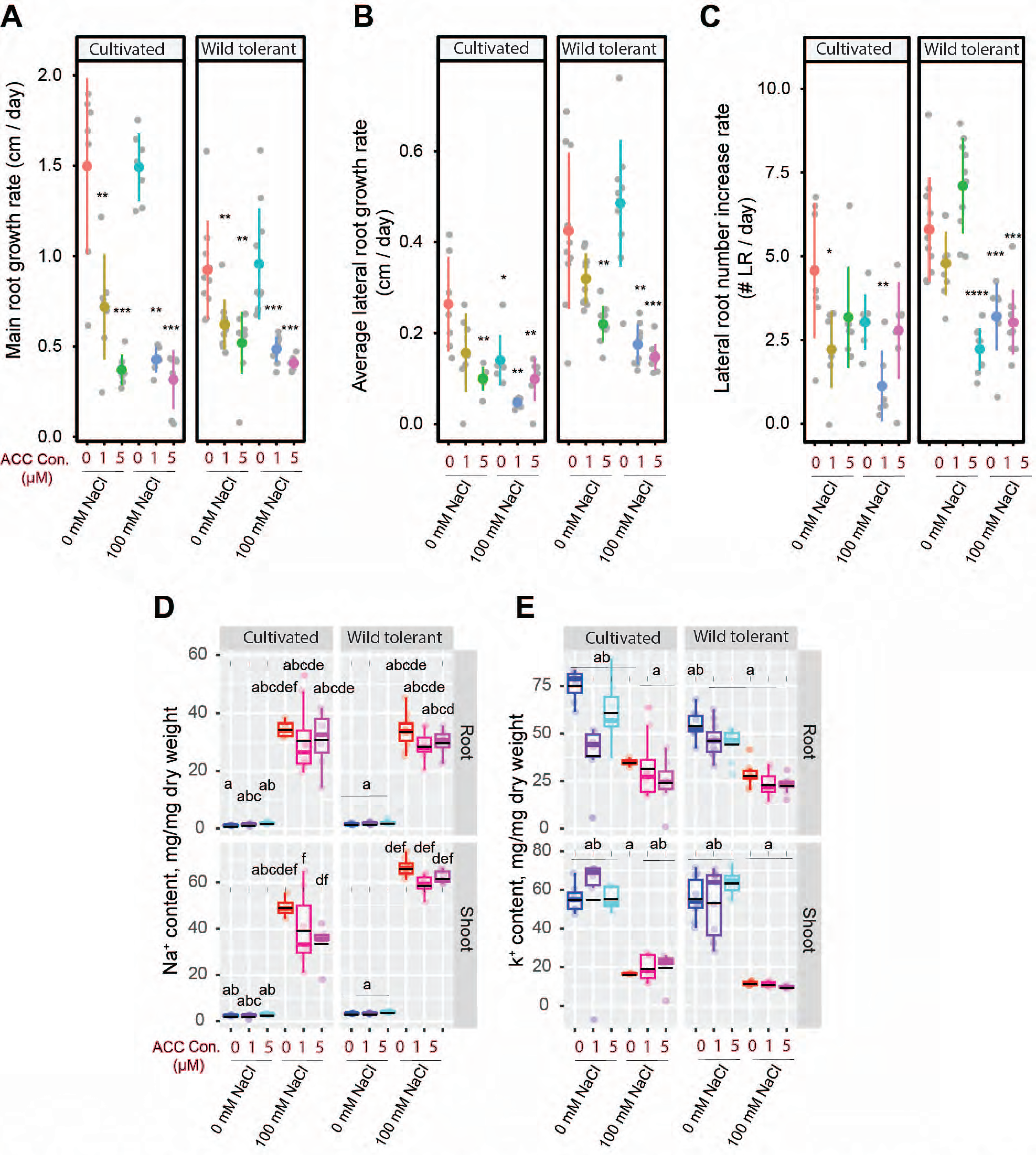


**Figure S13. ACC treatment significantly reduces both main root and lateral root lengths while causing a non-significant decrease in Na+ concentrations in the roots and shoots of both accessions under salt stress.** Root system architecture analysis of cultivated and wild tomatoes under 0 or 100 mM concentrations of NaCl supplemented with or without various concentrations of ACC, as indicated in the figure, are shown for main root growth rate **(A)**, average lateral root length **(B)** as well as average lateral root number **(C)**. Na+ **(D)** and K+ **(E)** content of root and shoot of different accessions after 10 days on treatment plates. (A-E) Each dot represents individual replicate per accession. (D-E) Lines represent mean values. The asterisks above the graphs in (A-C) indicate significant differences between control (i.e., 0 mM NaCl and 0 ACC) and other treatment by the Student 's t-test: *P < 0.05, **P < 0.01, ***P < 0.001, and ****P < 0.0001. Statistical analysis was done by comparison of the means for all pairs using Tukey–Kramer HSD test for (D-E). Levels not connected by the same letter are significantly different (P < 0.05).

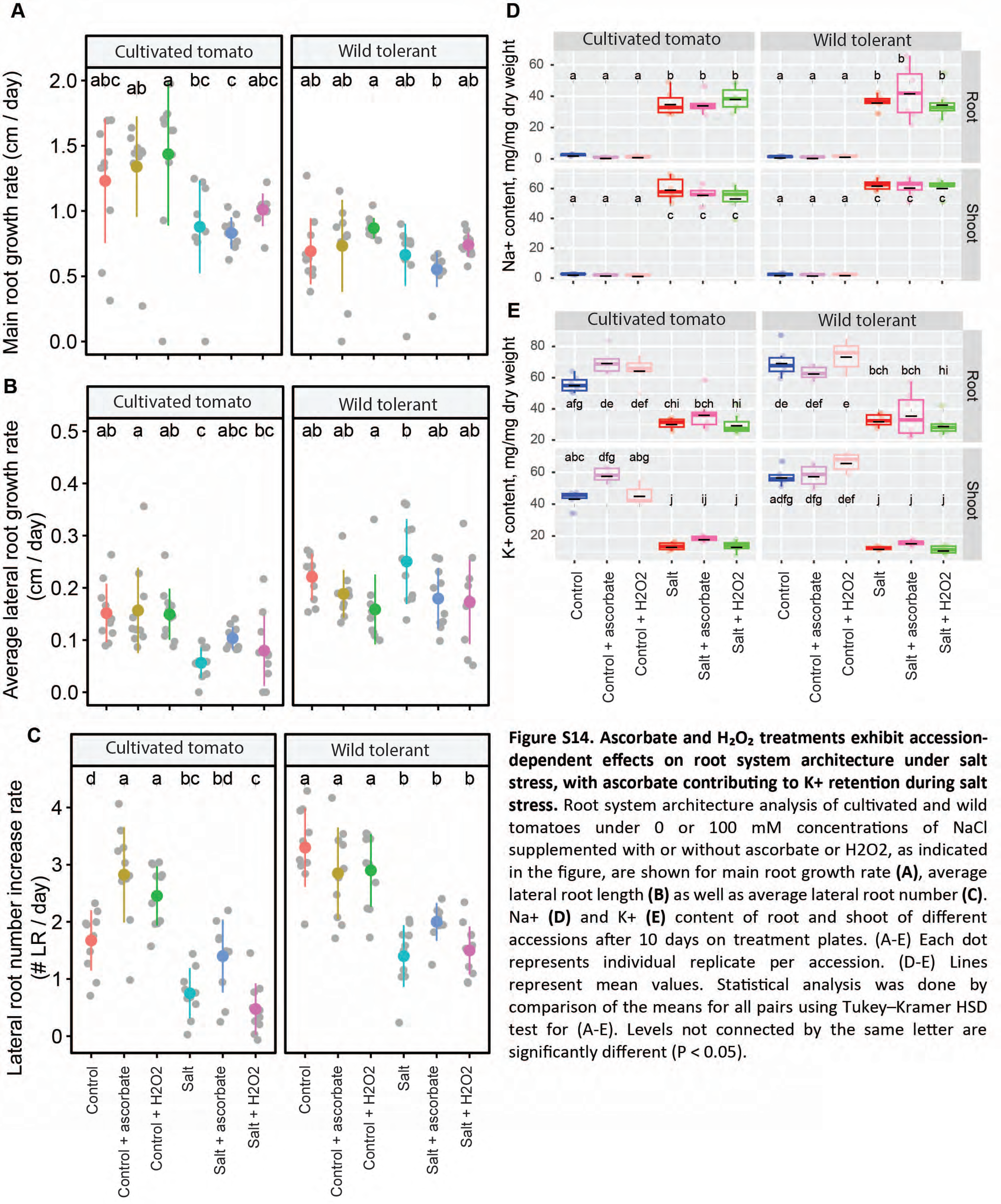

**Figure S14. Ascorbate and $H_2O_2$ treatments exhibit accession-dependent effects on root system architecture under salt stress, with ascorbate contributing to K+ retention during salt stress.** Root system architecture analysis of cultivated and wild tomatoes under 0 or 100 mM concentrations of NaCl supplemented with or without ascorbate or H2O2, as indicated in the figure, are shown for main root growth rate **(A)**, average lateral root length **(B)** as well as average lateral root number **(C)**. Na+ **(D)** and K+ **(E)** content of root and shoot of different accessions after 10 days on treatment plates. (A-E) Each dot represents individual replicate per accession. (D-E) Lines represent mean values. Statistical analysis was done by comparison of the means for all pairs using Tukey–Kramer HSD test for (A-E). Levels not connected by the same letter are significantly different ($P < 0.05$).
